# Genetic and functional analysis of unproductive splicing using LeafCutter2

**DOI:** 10.1101/2025.04.06.646893

**Authors:** Carlos F Buen Abad Najar, Ru Feng, Chao Dai, Benjamin Fair, Quinn Hauck, Jinghui Li, Xuewei Cao, Kushal K. Dey, Philip De Jager, David Bennett, The Alzheimer’s Disease Sequencing Project Functional Genomics Consortium, Xuanyao Liu, Gao Wang, Yang I Li

## Abstract

Alternative splicing commonly generates unproductive mRNA transcripts that harbor premature termination codons^1^, leading to their degradation by nonsense-mediated decay (NMD). These events reduce overall protein expression levels of affected genes, potentially contributing to gene regulation and disease mechanisms.

Here, we present LeafCutter2, which enables identification and quantification of unproductive splicing from short-read RNA-seq data. LeafCutter2 requires minimal gene annotations (start and stop codons) to annotate NMD-inducing splicing events, and identifies differential unproductive splicing between groups, providing insights into its contributions to differential gene expression. Moreover, LeafCutter2 enables mapping of unproductive splicing quantitative trait loci (u-sQTLs), which often colocalize with expression QTLs and GWAS loci.

Applying LeafCutter2 to RNA-seq data across human and 6 non-human species, we uncovered a broad landscape of unproductive splicing, which varies widely across tissues. Strikingly, we observed a conserved developmental-stage-specific increase in unproductive splicing during testis maturation across all species.

In Alzheimer’s disease (AD), we analyzed RNA-seq data from the AD Functional Genomics consortium FunGen-xQTL project and identified unproductive splicing events in 18 AD risk genes, including *TSPAN14*, *PICALM*, and *CASS4*, likely mediating genetic effects on disease risk. We performed an integrative analysis using gene expression QTLs, protein expression QTLs, and AD GWAS data, showing that unproductive splicing provides unique regulatory insights beyond traditional approaches. Thus, LeafCutter2 represents a powerful tool for understanding the functional impact of alternative splicing on gene expression and disease mechanisms.

## Introduction

Nearly all mammalian genes harbor introns that must be spliced out during mRNA maturation^2^. To achieve this, the spliceosome and auxiliary factors must bind pre-mRNA to 5’ and 3’ splice sites to initiate splicing^3,4^. Depending on the chosen splice sites, the introns that are removed can vary, producing alternatively spliced mRNA isoforms^4^. As the result of alternative splicing, a typical human gene can be transcribed and spliced into dozens of different mRNA isoforms. In some cases, these alternate transcripts encode different protein products^3–5^, but alternative splicing also frequently generates unproductive transcripts that contain a premature termination codon (PTC)^6–8^, leading to nonsense-mediated decay (NMD).

Because unproductive transcripts are lowly expressed in steady-state RNA-seq data^8^, they are difficult to detect and have been widely assumed to impact the regulation of just a small number of genes^9^. However, our recent work suggests a more significant functional role for this unproductive type of alternative splicing^1^. Indeed, we analyzed nascent RNA-seq from a human cell line to study transcript diversity prior to mRNA degradation and found that unproductive splicing is a global and potent negative regulator of steady-state mRNA expression levels^1^.

Additionally, we discovered that unproductive splicing often mediates the impact of non-coding variants on both gene expression level and disease susceptibility. These findings suggest that the coupling of alternative splicing and NMD (AS-NMD) is a major mechanism of gene regulation, and that the functional impact of alternative splicing events can often be interpreted in terms of whether they generate unproductive transcripts or not.

Yet there exists no streamlined method that can identify or quantify unproductive splicing events from RNA-seq data. Commonly used methods for splicing quantification such as MAJIQ^10^, rMATS^11^, Whippet^12^, and our original LeafCutter^13^ method, do not provide information on whether or not an alternative splicing event is expected to produce unproductive isoforms. This limits the functional interpretation of a huge number of mRNA transcripts.

Here we describe LeafCutter2, which allows the identification and quantification of unproductive splicing events from RNA-seq data. LeafCutter2 relies on minimal annotations – only the start and stop codons for each gene – and can thus be readily applied to RNA-seq data from human and non-human species with limited transcript annotations. We demonstrate LeafCutter2’s ability to quantify inter-tissue and inter-individual usage of unproductive splicing, which we confirm is associated with a concomitant difference in total mRNA expression levels. We apply LeafCutter2 to the vast amount of GTEx RNA-seq data from humans, as well as developmental RNA-seq data from 6 non-human species. We successfully identify tens of thousands of known and previously unannotated unproductive splicing events in humans and non-human species, revealing conserved usage of hundreds of unproductive splicing events that regulate gene expression. We further identify unproductive splicing QTLs (u-sQTLs) across 49 human tissues and identify colocalizations with complex trait GWAS loci. Finally, we apply LeafCutter2 to a large-scale RNA-seq dataset from Alzheimer’s disease (AD) brain samples and use state-of-the-art statistical genetics methods to uncover genes whose unproductive splicing likely influences AD risk. Overall, LeafCutter2 extends the functionality of the original package and substantially improves the functional interpretation of alternatively spliced isoforms.

## Results

### Overview of LeafCutter2

LeafCutter2 uses the same annotation-free approach to define clusters of alternative splicing as the original LeafCutter method^13^, leveraging split RNA-seq reads to identify intron excision events at base-pair resolution. This approach allows discovery of novel skipped exons, 5′ and 3′ alternative splice site usage, and additional complex events that can be summarized by differences in intron excision. The major advantage of this representation is that it does not require transcript reconstruction which is difficult and error-prone.

Briefly, LeafCutter pools mapped reads from all samples and finds overlapping introns specified by split reads. LeafCutter then constructs a graph connecting every overlapping intron that shares a donor or acceptor splice site. The connected components of the graph form clusters, which represent alternative intron excision events. Next, rarely-used introns are iteratively removed based on their usage relative to that of other introns in the same cluster, and new clusters are formed from the resulting disconnected subclusters. This last step is critical as it prevents the algorithm from clustering together the very large number of introns across the gene (this occurs often in large-scale RNA-seq data in which low-frequency, cryptic splicing events are commonly supported by multiple reads). Finally, because unproductive splicing events are often the product of cryptic splicing and/or are depleted in polyA RNA-seq data (as they are degraded by NMD), LeafCutter2 adds rarely-used junctions back to splicing clusters and recomputes relative usages (**Fig. 1a**).

**Figure 1:**
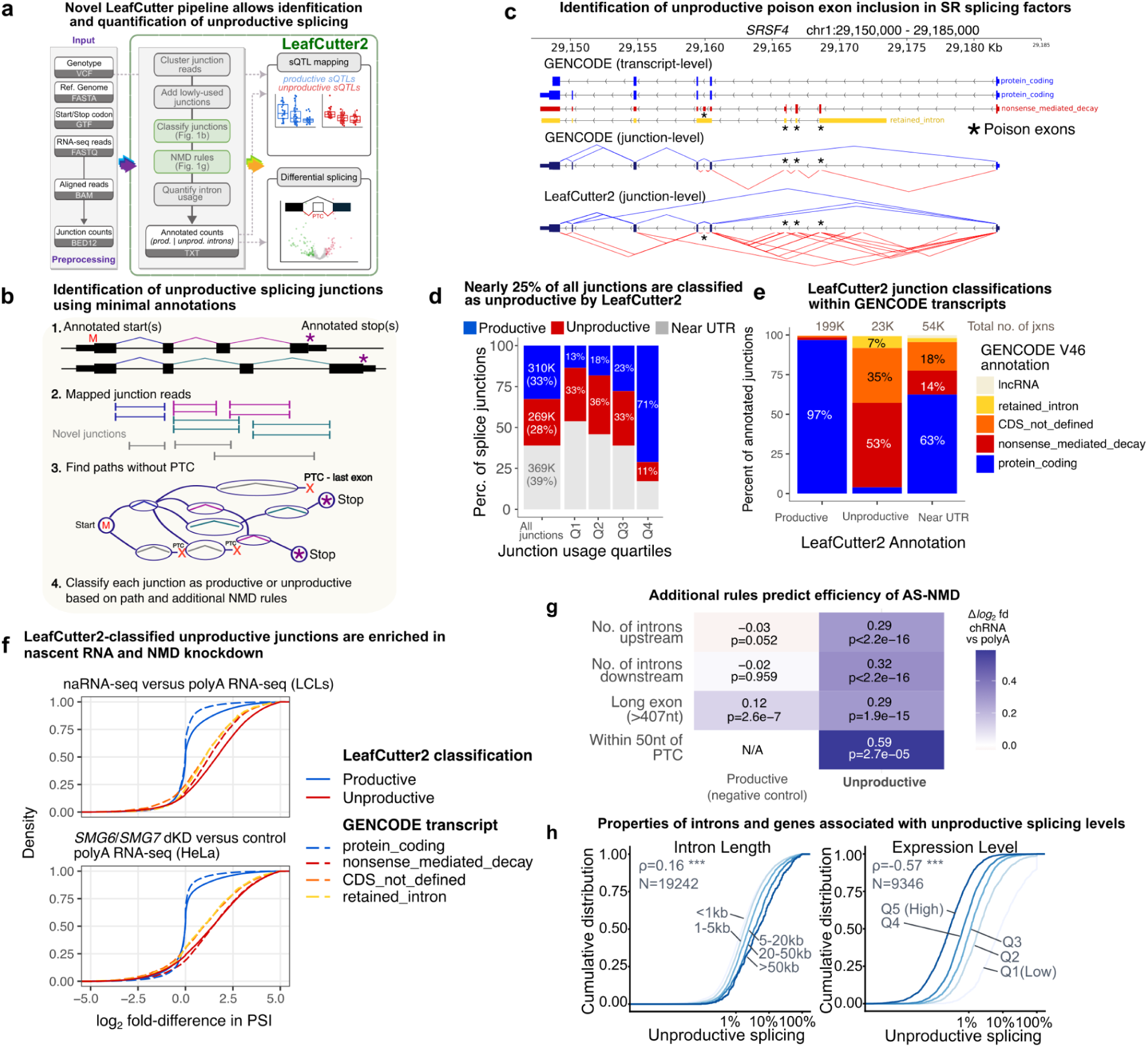
Robust annotation and quantification of unproductive splicing using LeafCutter2. **(a)** LeafCutter2 workflow. **(b)** Illustration of the breadth-first search approach to identify coding and NMD-inducing splice junctions. LeafCutter2 uses only start codon and stop codon as input to predict NMD-inducing junctions. **(c)** Example of annotated productive and unproductive splicing junctions in *SRSF4*. GENCODE annotations include junctions associated with three poison exons. LeafCutter2 annotates a large number of additional unproductive junctions. **(d)** Barplots showing the percentage of unique junctions annotated by LeafCutter as productive, unproductive, or near untranslated regions (UTRs). Percentage of all junctions shown, or junctions that are used at different levels (quartiles). The majority of highly-used (Q4) junctions are predicted to be productive, while only 14% are annotated productive for rarely-used (Q1) junctions. **(e)** Barplot showing GENCODE classification of junctions separated by LeafCutter2 annotations. Nearly all junctions annotated as productive by LeafCutter2 are annotated as belonging to protein coding transcripts in GENCODE. Unproductive junctions are nearly all exclusively within transcripts annotated as “nonsense_mediated_decay”, “CDS_not_defined”, or “retained_introns” in GENCODE. **(f)** Cumulative density plots showing the log2 fold-differences in Percent Spliced-In of junction reads between different types of RNA-seq libraries. Unproductive junctions classified by LeafCutter2 show enrichments (right-shift) in RNA-seq libraries expected to harbor an excess of NMD products. This level of enrichment is similar to that of junctions unique to GENCODE-annotated NMD transcripts. **(g)** Impact of additional rules that predict NMD efficiency. The 50nt rule is the strongest predictor of inefficient NMD. The number of introns upstream and downstream of an induced PTC is also an important predictor of NMD efficiency. **(h)** Cumulative density plots showing rates of unproductive splicing by intron length and host gene expression level. Longer introns and introns in lowly-expressed genes are most likely to exhibit high levels of unproductive splicing.

The main new feature in LeafCutter2 is the ability to identify splicing junctions with the highest probability of leading to an unproductive mRNA transcript, using minimal transcript annotations. LeafCutter2 uses a breadth-first search to consider all possible combinations of splice junctions and tries to find a path from an annotated start codon to a stop codon (GENCODE^14^ v46 annotation) without creating a premature termination codon (PTC). Specifically, LeafCutter2 employs a dynamic programming approach, leveraging the property that if there exists a PTC-free path from an annotated start codon to a specific 5’ splice site (5’ss), then all PTC-free paths from this 5’ss to an annotated stop codon will have a PTC-free path from an annotated start codon to an annotated stop codon (**Fig. 1b**). This allows LeafCutter2 to scale well even for long genes with hundreds of splicing junctions, taking less than an hour to annotate over 1 million splice junctions (**Supplementary Fig. 1**). We show an example of LeafCutter2’s annotations of productive and unproductive junctions in *SRSF4* (**Fig. 1c**), which nearly perfectly match with GENCODE v46 transcript annotations, though LeafCutter2 identifies many junctions not annotated by GENCODE.

In addition to identifying PTC relative to the annotated stop codon, LeafCutter2 implements rules posited to impact the efficiency of PTC to induce NMD: (i) the 50nt rule^15–17^, (ii) long exon rule^16,17^, (iii) total number of junctions upstream of the predicted PTC^17^, and (iv) total number of junctions downstream of the predicted PTC^17^ (**Fig. 1b**). These rules have been reported to influence the stability of mRNA transcripts with nonsense mutations^15–17^, but have not yet been systematically tested on unproductive mRNA transcripts resulting from alternative splicing. We show below that incorporating these rules improves LeafCutter2’s ability to predict the efficiency of NMD-mediated degradation of splicing junctions.

### Robust detection of NMD splice junctions using LeafCutter2

We evaluated LeafCutter2’s ability to identify unproductive splicing junctions by analyzing two complementary datasets: (i) RNA-seq data of double knockdowns of two core NMD factors (*SMG6* and *SMG7)* and shRNA controls from HeLa cell lines^18^ and (ii) nascent RNA-seq and steady-state RNA-seq data from lymphoblastoid cell lines (LCLs)^1,19^ (**Methods**). In total, LeafCutter2 extracted nearly 1 million unique splicing junctions from these datasets, including 269,331 (∼28% of all junctions) predicted to lead to unproductive transcripts. The remaining junctions were either predicted to be coding by LeafCutter2 (33%) or were too close to a potential untranslated terminal region (UTR) for us to accurately predict NMD status (39%).

When we examined junctions that are used at different frequencies, we found that the majority of frequently used splice junctions (71%) are predicted to be productive; however, this percentage decreases 13% when including splice junctions that are less frequently used (**Fig. 1d**). These observations are expected because most NMD-inducing junctions are likely results of low-frequency noisy splicing^1,20^. Still, we note that a substantial number of splice junctions annotated as unproductive are moderately or highly used (**Fig. 1d**), suggesting a functional impact.

When we focused only on the splice junctions annotated by both LeafCutter2 and GENCODE (v46), we found that 97% of "productive" junctions were located within protein-coding transcripts. A few exceptions involved junctions found only in NMD transcripts in GENCODE, but that LeafCutter2 found could still form protein-coding paths when combined with other detected junctions (**Supplementary Figure 2**). By contrast, splice junctions classified as unproductive were annotated within transcripts annotated as “nonsense_mediated_decay” (48%), “protein_coding_CDS_not_defined” (33%), or “retained_intron” (7%) (**Fig. 1e**). We observed that 3.7% of splice junctions classified as unproductive were annotated as protein-coding in GENCODE. Some of these junctions generate isoforms with a PTC within 50 nucleotides of an annotated stop codon, and are not expected to induce NMD as correctly assessed by GENCODE (**Supplementary Fig. 3**a). These can be reclassified based on additional NMD rules to improve annotation accuracy, as discussed below. Other junctions classified as NMD by LeafCutter2 correspond to transcripts that lack a defined open reading frame (ORF), but are nonetheless annotated as protein-coding by Gencode (**Supplementary Fig. 3**b). We conclude that, while GENCODE uses many sources of information for prediction in addition to manual curation, LeafCutter2 achieves near GENCODE-level annotations of unproductive transcripts with minimal gene annotations (just the start and stop codons).

To validate our classifications experimentally, we examined levels of unproductive splicing in samples subjected to NMD factor knockdowns, reasoning that unproductive junctions should be enriched in these samples relative to shRNA controls. We performed a similar comparison between nascent RNA-seq and steady-state RNA-seq, since unproductive transcripts should be more abundant at earlier time points, before NMD can break them down. As expected, we found that splice junctions classified as unproductive by LeafCutter2 were overwhelmingly enriched in RNA-seq of double *SMG6* and *SMG7* knockdowns^18^ and in nascent RNA-seq^1^ compared to shRNA control and polyA RNA-seq, respectively (**Fig. 1f**). LeafCutter2’s approach performs similarly compared to using the “nonsense_mediated_decay” transcript annotation from GENCODE (**Fig. 1f**), but increases the number of annotated unproductive splicing junctions by an order of magnitude (12.1-fold, **Supplementary Fig. 4**).

Using these data, we also evaluated multiple rules that previous studies have suggested impact the efficiency of a PTC to induce NMD based on its location^15–17^ (**Methods**). We found that all four implemented rules were useful predictors of a junction’s enrichment in nascent RNA-seq relative to steady-state RNA-seq. For example, comparing PTCs within long exons (>= 407nt) to those within short exons (<100nt), we observed a log_2_ fold-difference of 0.29 in nascent versus steady-state RNA-seq enrichment (**Fig. 1g**), suggesting that PTC within long exons are less effective at inducing NMD. By incorporating these added rules, LeafCutter2 accurately identifies splicing junctions leading to PTCs, and further distinguishes PTC-inducing junctions that are efficiently versus inefficiently degraded by NMD.

Finally, we examined the properties of introns or genes with low or high levels of unproductive splicing as quantified by LeafCutter2. We corroborated our previous findings^1^ that both long introns and low expression levels are significantly correlated with unproductive splicings **(Fig. 1h)**. Altogether, these analyses indicate that LeafCutter2 allows robust detection and quantification of NMD-inducing splice junctions from short-read RNA-seq data.

### Identification of unproductive splicing that result in tissue-specific gene expression

To evaluate LeafCutter2’s ability to detect unproductive splicing events in a large-scale RNA-seq dataset, we applied it to 17,329 GTEx samples spanning 49 tissue types (**Supplementary Table 1**). Notably, we observed that the proportion of unproductive splicing junctions present in sequenced reads varied significantly across tissue types, ranging from 0.5% in fibroblasts to 1.6% in testis (Fig. 2a), with a median of 1% of spliced reads classified as unproductive. We also found substantial variation across samples of the same tissue type. Some of this variation may be due to technical factors, such as differences in RNA integrity (**Supplementary Fig. 5**). Alternatively, it could reflect biological differences, such as certain cell types using higher levels of unproductive splicing to regulate gene expression. Another possibility is that the efficiency of degrading unproductive transcripts varies, leading to differences in the abundance of these transcripts in steady-state mRNA. While directly testing this hypothesis was beyond the scope of our study, we speculated that differences in NMD activity might explain the observed variation in unproductive splicing rates across tissues. To explore this, we analyzed the correlation between unproductive splicing levels and the mRNA expression of key NMD factors—*UPF1*, *UPF2*, *UPF3A*, and *UPF3B*—as proxies for NMD activity. Remarkably, we found a highly significant positive correlation (**rho =0.52, p < 1e-16**) between the expression level of *UPF3A* and unproductive splicing levels across tissue types (**Fig. 2a**) and across samples within tissue-types (**Supplementary Fig. 6**). Correlations with *UPF1, UPF2* and *UPF3B* expression levels across tissues were also significant (**Supplementary Fig. 7)**. Previous work on UPF3A suggests that it acts as a repressor of NMD in a mouse model and a human cell line^21^. Supporting this view, *UPF3A* expression levels in GTEx tissues strongly and positively correlates with unproductive splicing. Thus, our findings indicate that the efficiency of unproductive transcript degradation varies greatly across tissue types.

**Figure 2.**
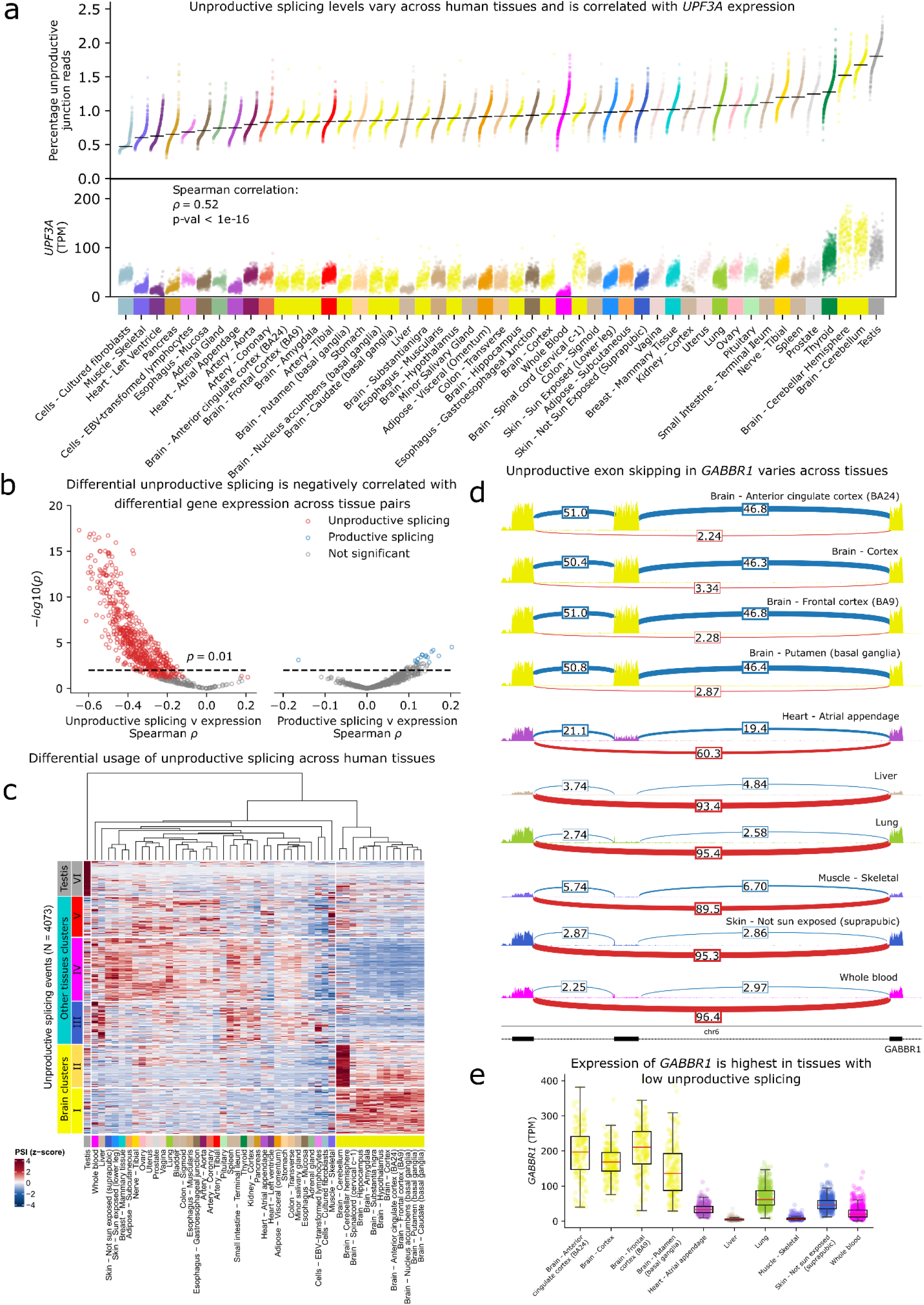
Differential unproductive splicing across human tissues. **(a)** Percent of junction reads that are classified as unproductive for each sample, grouped according to GTEx tissue type. **(b)** Scatter plot showing the Spearman’s correlation between delta PSI and differential gene expression Z-scores for tissue pairs (x-axis) and the significance of the correlation (y-axis). **(c)** Heatmap showing significant differential usage of unproductive splicing events (dPSI > 0.1) for genes that also show differential gene expression. **(d)** Sashimi plot of a differentially spliced unproductive event in *GABBR1* across GTEx samples. The unproductive event corresponds to an exon skipping event. **(e)** Boxplots of *GABBR1* expression level across GTEx tissues. High expression of *GABBR1* in brain tissue is associated with high exon inclusion.

Although NMD efficiency varies across cell types and influences the observed levels of unproductive splicing, it is likely that regulated unproductive splicing in specific genes can fine-tune their steady-state expression level in a tissue-specific manner. To investigate this possibility, we analyzed the differential usage of splicing junctions (difference in percent spliced in, or delta PSI) between tissue pairs for all junctions supported by at least 10 reads. We found a slight bias toward larger PSI for unproductive (but not productive) junctions in testis and cerebellum, consistent with reduced NMD activity in these tissues (**Supplementary Fig. 8**a,b). However, we also observed many unproductive junctions with substantially lower usage in testis or cerebellum compared to other tissues, indicating that at least some of the variation in unproductive splicing usage arise from regulated splicing rather than variation in NMD activity (**Supplementary Fig. 8**a,b). To further explore the potential regulatory significance of differential usage of unproductive splicing across tissue types, we examined the relationship between differential splicing of unproductive junctions and differential expression of impacted genes across tissue types. We used LeafCutter’s Dirichlet multinomial model^13^ to identify differentially spliced junctions across pairs of GTEx tissues, focusing on splicing clusters for which unproductive splicing junctions are estimated to have a differential splicing usage. This approach allowed us to adjust for the effect of NMD efficiency by using the percentage of unproductive splice junctions out of all splice junctions as a covariate (**Supplementary Fig. 8**c,d). At a 10% false discovery rate (FDR), we identified 5,289 differentially spliced unproductive events across one or more tissue-pairs with a minimum delta PSI of 10%. Of these, 4,596 events were in a gene that was also differentially expressed between the same tissues (10% FDR, minimum log_2_ fold change of 1). We then asked whether these differentially spliced unproductive events could explain cross-tissue differences in mRNA expression levels. To do so, we computed the correlation between the delta PSI of the unproductive junctions and the differential expression Z-scores of their host genes (**Methods**) for each tissue-pair separately. We expected that higher usage of an unproductive junction in a tissue would generally result in decreased mRNA expression levels. Indeed, we found that the correlations between delta PSI of unproductive junctions and differential gene expression Z-scores were overwhelmingly negative (635/765 pairwise comparisons with >50 differentially spliced unproductive clusters had a significant Spearman correlation between splicing and expression; 99.7% were negative) (**Fig. 2b, Supplementary Fig. 9**). This is not observed when using delta PSI of productive junctions (only 16 out of 1141 pairwise comparisons with >50 differentially spliced productive clusters had a significant correlation) (**Fig. 2b**).

To obtain a global view of tissue-specific gene regulation through unproductive splicing, we used hierarchical clustering to identify groups of unproductive splicing events that exhibit similar patterns of differential usage across tissue-types, restricting our analysis to the 4,073 unproductive splice junctions across 2,397 splicing clusters present in all 49 tissues to ensure robust clustering. We identified five notable patterns of tissue-regulated unproductive junction (**Fig. 2c**) with elevated usage in brain regions (group I), cerebellum (group II), liver/spleen (group III), skeletal muscle/heart and arteries (group V), and testis (group VI). Another large group of unproductive junctions were lowly used in brain regions and skeletal muscle/heart, but highly used in most other tissue-types (group IV, **Fig. 2c**). We show one illustrative example at the *GABBR* gene (**Fig. 2d**), where an unproductive junction is used predominantly in non-brain tissues, leading to lower mRNA expression levels in non-brain tissues relative to brain tissues.

Additional examples are shown in **Supplementary Fig. 10**. Thus, LeafCutter2 identifies thousands of unproductive splicing events that appear to tune the expression levels of their host genes across human tissue-types.

Finally, we conducted a gene ontology analysis to identify cellular functions that might be regulated post-transcriptionally, through the coupling of unproductive splicing and NMD. The most significant enrichments include genes involved in RNA splicing (OR = 2.15, adjusted p = 5.1×10^-4^ in gene group III, IV and V), response to DNA damage (OR = 2.14, adjusted p = 3.1 ×10^-7^ in gene group VI), and ncRNA metabolic processes (OR = 1.90, adjusted p = 3.8 ×10^-3^ in groups I and II), all of which are enriched in tissue-regulated unproductive splicing events relative to genes with at least one unproductive splicing junction but no tissue-specificity (**Supplementary Table 2**). These functional enrichments highlight a few specific examples of functional pathways whose basal activity is likely tuned by tissue-regulated unproductive splicing. However, thousands of genes with distinct functions harbor tissue-regulated unproductive splicing events, indicating that this mechanism likely impacts a diverse set of biological processes.

### Analysis of unproductive splicing in non-human species

To demonstrate the utility of LeafCutter2 for studying unproductive splicing in non-human organisms, we re-analyzed a large RNA-seq dataset encompassing various developmental stages across seven tissues in seven vertebrate species^22^. The human and mouse genomes have the most extensive gene annotations, often with more than 10 annotated transcripts for a single gene, many of which are unproductive (**Fig. 3a**). By contrast, macaque, rabbit, opossum and chicken have no annotated unproductive transcripts, with nearly all opossum genes having only one annotated productive transcript. This variability in annotation quality provided an opportunity to evaluate Leafcutter2’s accuracy in identifying NMD-inducing splice junctions in the absence of thoroughly annotated gene structures.

**Figure 3.**
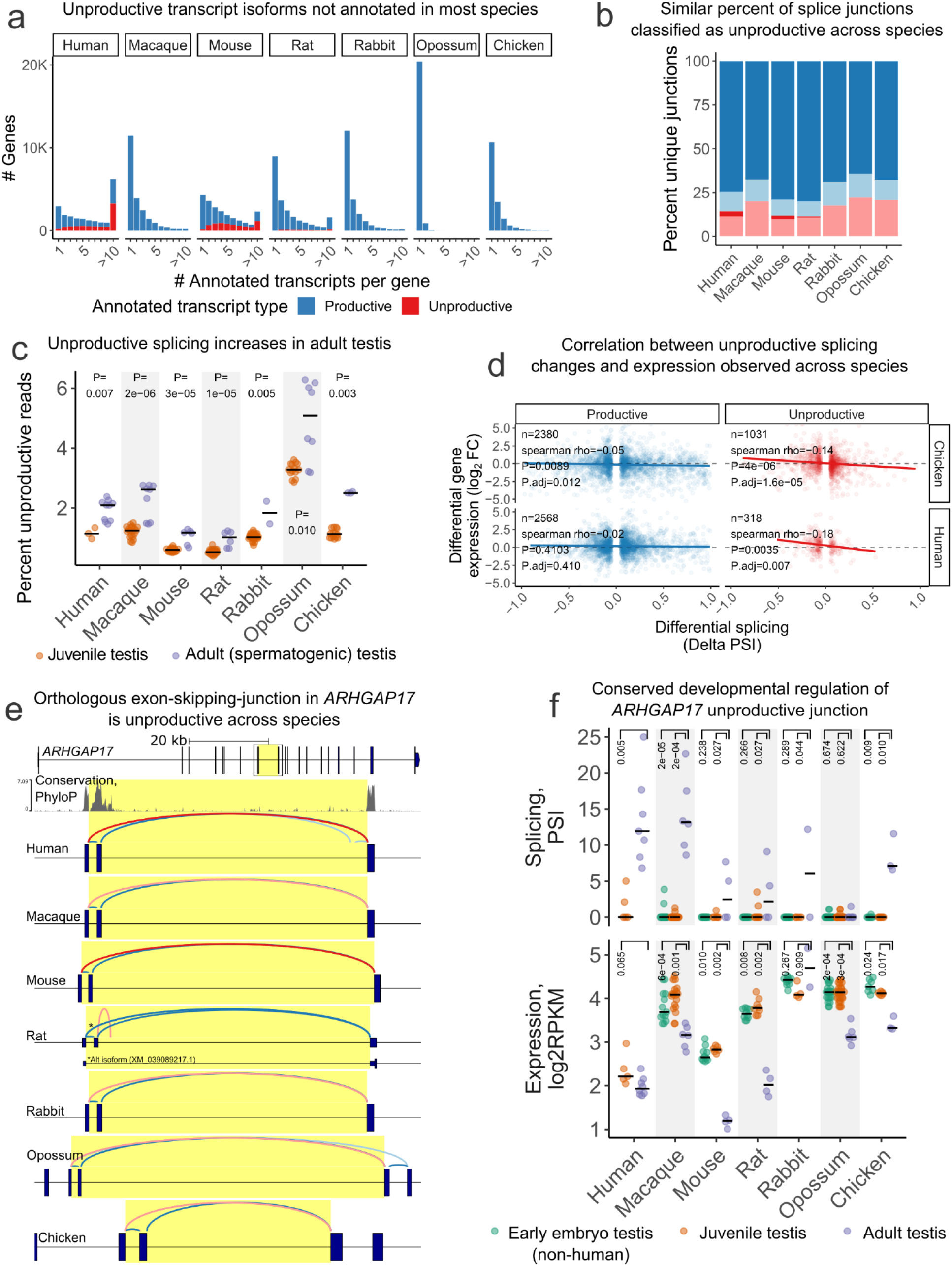
LeafCutter2 identifies unproductive splice events across species. (**a**) Histograms quantify the number of transcripts per protein-coding gene in each species’ current gene annotation. The fraction of transcripts that are productive are filled as a fraction within each bin. (**b**) Percent of unique junctions (filtering for junctions above a minimum abundance threshold, see Methods) that are productive (blue) or unproductive (red). Lighter shades represent junctions that are unannotated. (**c**) Percent of unproductive splice junction reads in each testis sample (point), comparing juvenile samples (defined as developmental stages between birth and onset of spermatogenesis) vs adult (spermatogenic). Group medians are represented with dashes. The p-values shown are from a two-sided Mann Whitney test. (**d**) Scatter plots depict the correlation between differential splicing effect sizes over human and chicken testis development and host-gene differential expression effect sizes for each significant differentially spliced junction (FDR < 1%, DeltaPSI > 5%). Splice junctions in LeafCutter2 clusters with a significantly differentially spliced unproductive junction (“Unproductive”) are evaluated separately from those without (“Productive”). Correlation summarized with rho coefficient and the p-values shown correspond to two-sided Spearman correlation tests. (**e**) Full productive human gene structure of *ARHGAP17* is shown as the top track, with the box showing zoomed in region on the tracks below. Gene structure of orthologous region is shown for each species, with orthologous junction highlighted. All splice junctions in LeafCutter2 cluster are drawn as arcs, colored by productive/unproductive status and annotation status as in (**a**). The 1000 vertebrate nucleotide conservation PhyloP track is plotted above the human track. *The rat orthologous junction is an annotated productive junction, due to an annotated isoform with an alternative transcription start site, though the junction is predicted to be NMD-inducing in the context of the main, full-length transcript. (**f**) (top) Splicing of junction highlighted in (**e**), and (bottom) *ARHGAP17* expression levels for non-human early embryo, juvenile, and adult testis samples are shown as points for individual samples, summarized as black dash for median within each group. The p-values shown are from a two-sided Mann-Whitney test.

We aligned RNA-seq reads to their respective genomes, extracted splice junction reads, and used LeafCutter2 to identify the splice junctions most likely to be specific to unproductive transcripts (**Methods**). We extracted a similar number of unique splice junctions in each species (ranging from 0.76 to 1.22 million junctions). Of these, about 30% (range of 27.5 – 33.8) were present at an appreciable abundance, with approximately 50 junction reads or greater (**Methods**). Leafcutter2 classifies a similar proportion of these abundantly expressed junctions as unproductive across the seven species (12-22%, **Fig. 3b**), despite the vast differences in annotation quality. When considering the fraction of splice junction *reads* that are unproductive in each sample, similar to our analysis of GTEx samples (**Fig. 2a**), we find that most samples contain 0.5-2% unproductive junction reads, with some variability between species, tissues, and developmental stages (**Supplementary Fig. 11**). These measures of the total fraction of unproductive splice junction reads are similar whether using annotations from various sources (e.g., RefSeq, Ensembl) which sometimes differ in comprehensiveness even within species (**Supplementary Fig. 12**). Consistent with the elevated levels of unproductive splicing we identified in adult testis samples from GTEx (**Fig. 2a**), we find adult testis samples exhibit elevated unproductive splicing across all seven species in this dataset (**Fig. 3c**). This elevation in unproductive splicing coincides developmentally with sexual maturation of the testis (**Fig. 3c**), as well as an increase in the expression of *UPF3A* relative to *UPF3B* expression in most species (**Supplementary Fig. 13**). While this increase in *UPF3A*/*UPF3B* is partly or fully due to an increase in *UPF3A* expression in some species (e.g. chicken), in other species it is partially or fully driven by a decrease in *UPF3B* expression, likely due to mammalian X-chromosome inactivation of *UPF3B* during testis maturation^23^. This is consistent with UPF3A’s role as antagonist of *UPF3B* and an inhibitor of NMD^21^, and suggests that maintaining the appropriate balance of UPF3A/UPF3B and global partial inhibition of NMD during testis maturation is likely conserved across amniotes, and can be achieved in multiple ways.

Having classified splice junctions as unproductive or productive across species with less-comprehensive annotations, we next sought to verify the accuracy of these classifications. We looked for the same anti-correlation between splicing of unproductive junctions and the expression of host-genes that we observed in GTEx data (**Fig. 2c**). Specifically, we identified significant splicing changes between developmental stages and tissues within each species and compared those developmental splicing changes to the corresponding expression changes in host genes (**Methods**). For example, we find 318 significant unproductive and 2,566 significant productive splicing changes when comparing human kidney to brain (**Fig. 3d**). While the deltaPSI of the productive splicing changes does not strongly correlate with host-gene expression changes, the unproductive splicing changes are strongly negatively correlated, as expected (**Fig. 3d**). We obtained similar results in the analogous analysis in chicken tissues where the productive/unproductive classifications were made with a less comprehensively annotated genome. While differences in sample sizes impaired our power to assess the correlation between unproductive splicing and gene expression in some tissues and developmental stages, the same negative correlation was generally recapitulated in well-powered comparisons across all species (**Supplementary Fig. 14**). Overall, these results suggest that LeafCutter2’s classifications are accurate even in species with less comprehensive annotations.

Notably, the negative correlation between unproductive splicing changes and host gene expression is observed when comparing juvenile to adult (spermatogenic) testis (**Supplementary Fig. 14**). If the increases in unproductive splicing during testis maturation were solely due to a global inhibition of NMD, we would instead expect a slight positive correlation - inhibition of NMD would increase unproductive splicing and host gene expression.The observed negative correlation supports the idea that many of the increases in unproductive splicing during testis maturation are driven by changes in splicing, not just relief of NMD, and this contributes to down-regulation of host genes. Given this relationship, we next sought to identify unproductive splicing events that are conserved across species, as their conservation suggests regulatory importance.

We identified thousands of conserved splicing events using progressively stringent filters for evidence of conservation (**Methods**, **Supplementary Fig. 15**a). Among the 7,028 unproductive alternative splicing events we identified in humans and at least one other non-primate species, we recovered many known examples of conserved cassette exons (**Supplementary data file 4**) that are unproductive when included (“poison exons”), such as the fetal- and non-brain-isoform of *SCN8A* (**Supplementary Fig. 15**b-e) and the auto-regulatory poison exons in SR-family splice factors (e.g. *SRSF3*, *SRSF4*, *SRSF6*) (**Supplementary Fig. 15**e). Many of these poison exons are highly conserved at the nucleotide level while lacking signatures of strong amino acid coding constraint (**Supplementary Fig. 15**d-e), consistent with selection targeting the regulatory potential of the alternative splicing event. In contrast, constitutive exons flanking these unproductive cassette exons and productive cassette exons do not show the same signature of nucleotide conservation without amino acid constraint. Notably, some of these conserved poison exons are in genes unrelated to RNA biology (e.g., the cell-cycle regulator *CCNL1*, the cytoskeleton gene *ACTN1*, and the stress-response transcription factor *NFAT5*), suggesting AS-NMD may be a widespread mechanism regulating diverse biological processes.

Finally, we highlight an AS-NMD event in the rho-GTPAse activating gene *ARHGAP17* (**Fig. 3e**), whose regulatory function has not been previously described to our knowledge. This event is one of the 630 AS-NMD events we identified as being tissue- or developmentally-regulated in a conserved manner (**Supplementary Fig. 15**a). The increase of this unproductive exon-skipping junction from juvenile to adult testis is significant in all species examined except opossum (**Fig. 3f**). To assess whether the increase in this *ARHGAP17* unproductive splice isoform is merely a reflection of partial NMD inhibition, or a genuine increase in *ARHGAP17* unproductive splicing, we assessed how *ARHGAP17* expression level changes. Consistent with an increase in unproductive splicing and the regulatory potential of this event, expression of *ARHGAP17* from juvenile to adult testis is decreased in most examined species (**Fig. 3e**). Because this splice event generates an unproductive transcript by exon skipping rather than poison exon inclusion, the cassette exon is not expected to show the signature of nucleotide-but-not-amino acid constraint. However, we observe extended nucleotide conservation of the intronic regions flanking the skipped exon (**Fig. 3e**, **Supplementary Fig. 15**f), consistent with the functional importance of this unproductive splice event. Taken together, our results show that LeafCutter2 can be used to annotate and quantify unproductive splicing in non-human species, revealing an underappreciated preponderance of conserved unproductive splicing events with developmental- or tissue-specific patterns.

### Unproductive splicing mediates the effect of genetic variants on gene expression across human tissues

We previously collected nascent RNA-seq from a panel of 86 LCLs and used these data to map nearly one thousand genetic variants associated with unproductive splicing, i.e. unproductive splicing QTLs, or u-sQTLs^1^. We found that u-sQTLs colocalized with GWAS loci at a rate similar to sQTLs that impact productive splicing events, indicating that trait-associated variants often exert their impact through NMD-induced changes in gene expression levels^1^. Here, we used LeafCutter2 to identify u-sQTLs using polyA RNA-seq data from GTEx across the 49 available GTEx tissues.

We define p-sQTLs as genetic loci affecting intron clusters that exclusively lead to productive mRNA isoforms as annotated by LeafCutter2, and u-sQTLs as variants affecting mixed clusters that can lead to both productive and unproductive isoforms. We mapped thousands of significant p-sQTLs (32,334 unique clusters across all tissues, median 3,082 clusters per tissue) and u-sQTLs (16,816 unique clusters across all tissues, median 1,800 clusters per tissue) across all 49 tissues (**Fig. 4a**). These p-sQTLs and u-sQTLs affect alternative splicing of 9,387 and 5,107 genes, respectively, in at least one tissue. Their summary statistics are available online, formatted for easy overlap and statistical genetics analyses (Zenodo DOI: <u>10.5281/zenodo.15098365</u>). As expected, the number of p-sQTLs and u-sQTLs discovered per tissue is strongly correlated with the number of genotyped samples per tissue (Spearman rho: 0.92, p < 1e-16). Additionally, we note that we mapped many fewer u-sQTLs compared to p-sQTLs, which likely results from the depletion of unproductive splice junction reads in polyA RNA-seq^1^.

**Figure 4.**
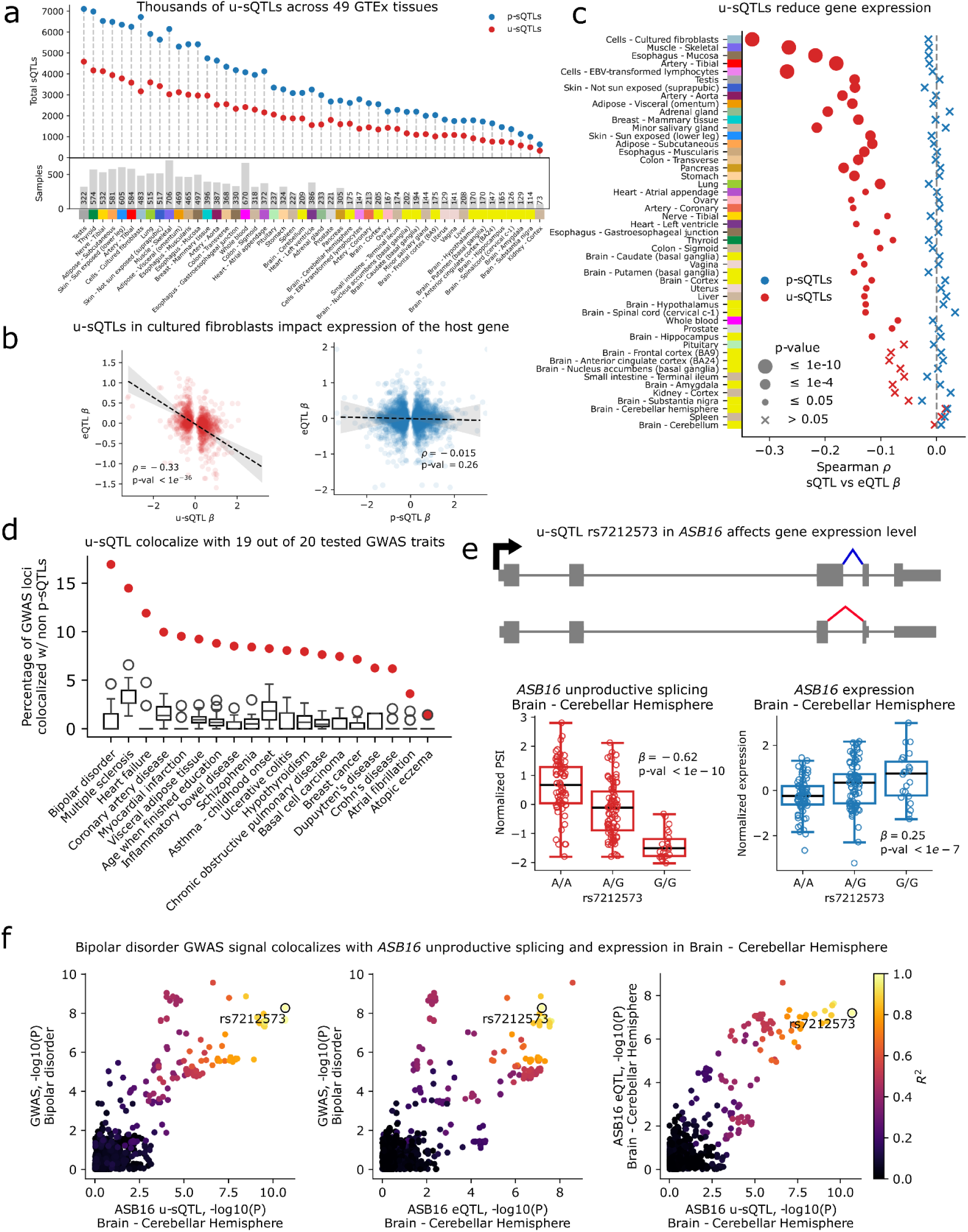
Genetic basis of variation in unproductive splicing. **(a)** Scatter plot showing the number of productive and unproductive sQTLs identified across GTEx tissues at 10% False Discovery Rate. **(b)** eQTL vs u-sQTL and p-sQTL effect size comparisons for an example tissue (cultured fibroblast). Alleles that increase unproductive splicing – but not productive splicing – tend to decrease expression level of the host genes. **(c)** Summary of comparisons between eQTL and sQTL effect sizes across all GTEx tissues, separating u-sQTLs and p-sQTL as classified by LeafCutter2. (**d**) Boxplots showing the percentage of GWAS colocalized with a u-sQTL identified in GTEx tissues. Red dot donates the total number of unique loci colocalized with at least one u-sQTL in any tissue. No colocalizations were found between u-sQTLs and any rheumatoid arthritis loci. (**e**) Example of a u-sQTL that colocalizes with bipolar disorder. The allele associated with increased bipolar disorder risk increases unproductive splicing in *ASB16*, and decreases its expression levels. (**f**) LocusCompare plots showing the colocalization of the GWAS association, eQTL, and u-sQTL at *ASB16*.

Since unproductive splicing is predicted to result in mRNA degradation through NMD, we verified that alleles that increase unproductive splicing have strong, negative effects on host gene expression levels, as illustrated for u-sQTLs identified in fibroblasts (**Fig. 4b**). By contrast, the effects of variants affecting p-sQTLs are not significantly correlated with gene expression levels (**Fig. 4b**). The negative correlation between genetic effects on unproductive splicing levels and gene expression levels is significant in the majority of the 49 tissue types we analyzed, with the strength and significance of the association being strongly correlated with the number of significant u-sQTLs found per tissue (**Fig. 4c**). By comparison, p-sQTLs show no such negative correlation in any tissue. Consistent with these results, we found that u-sQTLs have a stronger eQTL effect for their host genes than p-sQTLs across all 49 tissues **(Supplementary Fig. 16)**.

Next, we tested for colocalization between u-sQTLs and a total of 2,897 GWAS loci from 20 complex traits (**Supplementary Table 3**). We found that 225 GWAS loci, or 7.8%, colocalize with at least one u-sQTL in at least one GTEx tissue (**Fig. 4d**). One example case is rs7212573, an intronic variant in *ASB16*, a gene of the ankyrin repeat and SOCS box-containing family that is involved in protein metabolism. The minor allele at rs7212573 is associated with increased unproductive splicing of *ASB16*, which results in decreased expression of *ASB16* (**Fig. 4e,f**) and an increase in risk for bipolar disorder. We include two additional examples where unproductive splicing likely explains the impact of a hyperthyroid-associated locus and a breast cancer-associated locus on the expression levels of *IRF7* and *PIDD3*, respectively (**Supplementary Fig. 17, Supplementary Fig. 18**). These observations highlight the widespread impact of genetic variants affecting unproductive splicing on gene expression across human tissues.

### Genetic basis of unproductive splicing reveals novel regulatory mechanisms in Alzheimer’s disease

We have shown that LeafCutter2 can accurately identify unproductive splicing events from minimally annotated transcriptomic data. By applying LeafCutter2 to RNA-seq datasets in multiple tissues and across species, we found evidence that AS-NMD is a widespread and evolutionarily conserved mechanism of tissue-specific gene regulation. We next investigated the utility of this approach for investigating complex disease, specifically Alzheimer’s disease (AD), hypothesizing that unproductive splicing and its genetic regulation could contribute to the molecular mechanisms underlying established AD susceptibility loci. Specifically, we used LeafCutter2 to analyze data from the Religious Orders Study and Memory and Aging Project (ROSMAP), a substantially larger brain resource compared to GTEx, enabling us to establish a comprehensive mapping of unproductive splicing events and u-sQTLs in the aging human brain. In the ROSMAP dataset, we mapped splicing QTLs, including both u-sQTLs and p-sQTLs, in three brain regions: dorsolateral prefrontal cortex (DLPFC, n=806), anterior cingulate cortex (AC, n=603), and posterior cingulate cortex (PCC, n=449). For DLPFC, we further integrated these splicing QTLs with additional molecular QTLs generated by the ADSP Functional Genomics Consortium xQTL Project (FunGen-xQTL), including expression QTLs from both bulk tissue and single-nucleus RNA-seq, protein abundance QTLs (pQTLs), and H3K9 histone acetylation QTLs (haQTLs). Key details of these molecular QTL datasets are outlined in **Methods**, with more comprehensive information provided in the FunGen-xQTL manuscript^24^.

We used LeafCutter2 to identify 364,225, 351,485 and 338,633 introns grouped into 108,420, 120,988 and 106,399 intron clusters from RNA-seq data of DLPFC, AC, and PCC, respectively. Through QTL association analysis, we mapped 3,609, 4,517, and 3,459 u-sQTLs at a 10% FDR in the DLPFC, AC, and PCC, respectively, as well as 6,026, 8,071, and 6,346 p-sQTLs at 10% FDR across these brain regions. In the DLPFC, we further investigated how both types of splicing QTLs relate to other molecular regulatory mechanisms by comparing them with eQTLs, pQTLs, and haQTLs. Consistent with our GTEx analysis, u-sQTLs showed stronger enrichment for eQTL signals compared to p-sQTLs, suggesting that u-sQTLs are more likely to influence gene expression levels. A similar enrichment pattern was observed with pQTLs (**Fig. 5a**). In contrast, we found no enrichment with haQTLs, which regulate chromatin accessibility through histone modifications prior to transcription (**Supplementary Fig. 19**). This absence of correlation aligns with the post-transcriptional nature of NMD regulated by u-sQTLs.

**Figure 5.**
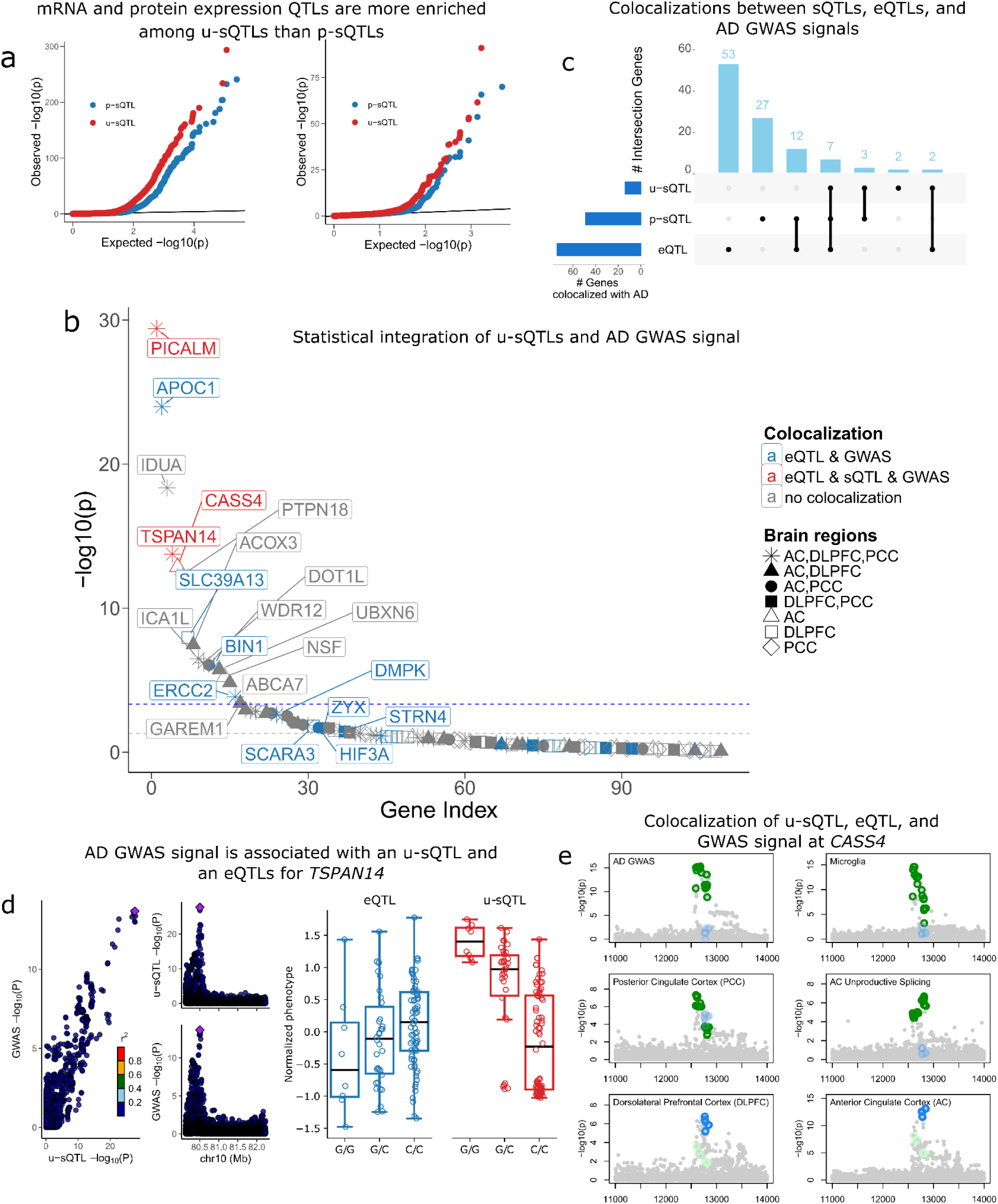
Impact of unproductive splicing on Alzheimer’s disease. **(a)** Quantile-quantile (QQ) plots showing DLPFC eQTL (left) and protein expression QTL (right) signal among u-sQTLs and p-sQTLs. As expected, u-sQTLs show stronger enrichment among eQTLs and pQTLs as compared to their productive counterparts. **(b)** Scatter plot showing the integration of u-sQTL effects with AD GWAS through TWAS (y-axis) and colocalization (colored legend) analysis. Several signals were identified using u-sQTLs from multiple brain regions. The gray dashed line represents p = 0.05; the blue line indicates the multiple testing-corrected p-value threshold, adjusted based on the number of tests performed. Significant colocalization was defined by PP.H_4_ > 0.8. **(c)** UpSet plot showing results of a multi-trait colocalization across u-sQTLs, p-sQTLs, eQTLs, and AD GWAS. **(d)** Regulatory effects at the *TSPAN14* locus. Left: LocusCompare plot showing colocalization between AD GWAS and u-sQTL signals. Right: Boxplot showing the opposite genetic effect on unproductive splicing and *TSPAN14* expression level. **(e)** Manhattan plot showing multi-context genetic effects at the *CASS4* locus. Green points indicate variants colocalized between AD GWAS, *CASS4* bulk eQTL. (DLPFC, AC, PCC), sc-eQTL from microglial, and u-sQTL for a splicing cluster in *CASS4*. Blue points show a separate colocalization signal specific to DLPFC and AC *CASS4* eQTL. To illustrate their relationship, we overlay light green points on the secondary signal to show that primary colocalization variants are only moderately significant, and light blue points on the primary signal to indicate a similar moderate significance for the secondary variants.

Using DLPFC data, we investigated how u-sQTLs contribute to AD risk by performing an in-depth analysis of 244 candidate genes generated through a multi-step prioritization process that integrated multi-context molecular QTL evidence, fine-mapped AD GWAS meta-analysis, and complementary functional genomics data (e.g. ClinGen, LDH, and computational prioritization tools)^24^ (**Supplementary Table S4**). To achieve robust inference, we employed multiple QTL integration approaches on summary statistics from a recent AD GWAS meta-analysis^25^. First, we performed a probabilistic transcriptome-wide association study (PTWAS)^26^ to identify AD-relevant genes driven by u-sQTL. We also performed pairwise colocalization using SuSiE-COLOC^27^ to find shared putative causal variants between u-sQTL and AD GWAS. To mitigate possible TWAS false positives due to linkage disequilibrium hitchhiking^28^, we cross-referenced PTWAS identified AD risk genes (Bonferroni-adjusted p-value < 0.05) with independent evidence from pairwise colocalization between splicing/expression QTL and AD GWAS data.

Among 17 risk genes identified through TWAS with unproductive splicing, three genes — *PICALM*, *TSPAN14* and *CASS4* — also exhibited strong evidence of colocalization between u-sQTL and GWAS, as well as colocalization with at least one eQTL context (**Fig. 5b, Supplementary Table S5**). To systematically investigate whether u-sQTL signals converge with other layers of molecular regulation, we further applied ColocBoost^29^ — a powerful multi-trait colocalization method developed by the FunGen-xQTL consortium^24^ — to jointly analyze sQTL with other multi-context eQTL data. Our analysis revealed 7 genes for which genetic variants may regulate both unproductive splicing and gene expression levels, reinforcing the notion that these variants contribute to AD risk through coordinated regulatory cascades (**Fig. 5c**).

To demonstrate how our analyses may reveal novel insights into AD pathology, we examined several prioritized genes in detail. At the *TSPAN14* locus, colocalized genetic effects were consistently observed across all three brain regions, providing robust evidence for a splicing-mediated mechanism. This finding is supported by recent experimental validation using long-read sequencing, which confirmed splicing regulation of *TSPAN14* as an established AD risk factor^25^. By integrating multi-context colocalization with cell-type-specific expression data, we further discovered that these genetic effects were particularly pronounced in excitatory neurons (**Supplementary Fig. 20**a), suggesting a specific cellular context for the role of *TSPAN14* in AD pathogenesis. While mediation analysis confirmed that splicing changes significantly affect gene expression levels of *TSPAN14* (FDR < 0.05, **Methods**), the proportion of expression variation explained was modest, indicating that additional regulatory mechanisms beyond splicing contribute to expression variation of this gene (**Fig. 5d, Supplementary Fig. 20**b,c).

Our analysis also revealed previously unreported regulatory effects at known AD risk loci. For example, at *CASS4*, the most recently identified member of the CAS protein family involved in tyrosine kinase binding^30^, we observed complex splicing regulation that may explain its role in AD risk. Through multi-context colocalization, we identified shared genetic effects between AD GWAS, DLPFC microglial expression, PCC bulk expression, and unproductive splicing in AC (**Fig. 5e**). Notably, *CASS4* showed microglia-specific expression in our single-nucleus data, suggesting potential immune-related functions that have not been previously characterized. We further detected a secondary eQTL colocalization signal in DLPFC and AC close to but not overlapping with the u-sQTL. Given the moderate linkage disequilibrium between these loci (r = 0.12, **Methods**), they likely represent distinct causal variants with different functional effects. While CASS4 did not exhibit significant mediation of unproductive splicing effects on either bulk or microglial gene expression, the observed colocalization between u-sQTLs and AD-associated variants suggests a potential regulatory role of NMD at CASS4, particularly in microglia, that may contribute to AD risk and warrants further investigation.

Similar patterns of coordinated molecular regulation were observed at several other AD loci. At the established AD risk gene *PICALM*, we identified unproductive splicing effects colocalized with fine-mapped AD susceptibility variants, suggesting a possible uncharacterized transcript-level regulation. At *CTSH*, we observed strong colocalization between u-sQTLs and AD GWAS variants within an intronic region, supporting a potential functional role for *CTSH* in AD risk despite only suggestive AD association based on GWAS data alone (p-value = 5.9e-06)^25^. These examples demonstrate how integrating u-sQTLs with genetic and molecular data can expand our understanding of both well-established and emerging disease risk factors. Additional loci and u-sQTL regulatory patterns are explored in depth in other companion FunGen-xQTL consortium manuscripts^24,29^.

## Discussion

Most prior studies have focused on the role of alternative splicing (AS) in generating alternative protein isoforms, which can have important functional roles but are challenging to characterize (see references^31,32^ for notable efforts). However, a significant proportion of alternative splicing events are not expected to generate mRNA isoforms that encode proteins with alternative function, as they introduce premature termination codons that trigger NMD rather than being translated. Capturing these isoforms can be difficult since they are rapidly degraded following transcription and thus are present at relatively low abundance.

Here, we designed LeafCutter2, a streamlined pipeline that can predict whether or not splice junctions will lead to premature termination codons. This allowed us to predict the functional consequence of a large number of alternative splicing events. Indeed, at least 28% of all human splice junctions detected in this study lead to unproductive transcripts which result in rapid degradation by NMD. Even when considering junctions that are used at appreciable levels (above median usage), about 22% of all junctions (∼105k total junctions) are predicted to be unproductive. Thus, LeafCutter2 helps interpret the functional impact of up to a hundred thousand well-expressed human mRNA isoforms.

Our analysis revealed that the levels of unproductive splicing detected in polyA RNA-seq datasets vary significantly across different tissue types and show a strong positive correlation with *UPF3A* expression levels. This suggests that the efficiency of degrading unproductive transcripts through the NMD pathway is regulated differently across cell types, though the precise function of this regulation – if any – remains unclear. Additionally, we identified thousands of unproductive splicing events that are differentially regulated across tissues, even after accounting for variation in NMD activity. Among these, hundreds are conserved across multiple species and likely play a role in fine-tuning gene expression in a functional and tissue-specific manner. Many more such events likely exist but are challenging to detect using steady-state polyA RNA-seq data.

Our work highlights the potential value of creating more comprehensive maps of unproductive splicing events across cell types and species. Approaches such as NMD factor knockdowns^18^ or sequencing of chromatin-associated RNA^1^, particularly when combined with long-read sequencing, could help distinguish between tissue-specific splicing regulation and variation in NMD activity. In the context of evolutionary biology, maps of unproductive splicing events prior to NMD could identify conserved unproductive splicing events with functional significance. In the context of human genetics, these maps could provide additional insights into the effects of genetic variants on alternative splicing and gene expression.

Analysis of unproductive splicing also offers the potential to uncover insights into the causal mechanisms of various complex diseases. Although we focused on using LeafCutter2 to study genetic loci associated with AD as a flagship example, emerging evidence suggests that AS-NMD may play a broader role in neurological disorders and immune functions, such as in ALS^33^ and viral responses^34^. Further research will likely reveal many more types of disease impacted by this mechanism, given its apparent regulatory role in a wide variety of tissue types. Despite limited systematic investigation of this mechanism in previous large-scale QTL and GWAS studies, our development of Leafcutter2 now enables high-resolution, annotation-free discovery of unproductive splicing, creating new opportunities to leverage GWAS data in a more mechanistic way.

## Online Methods

### LeafCutter2 classifies splice junctions in a computationally efficient manner

The main innovation of LeafCutter2 is a dynamic programming algorithm that classifies all splice junctions as productive if they can form a valid protein-coding transcript, or unproductive if they introduce a premature termination codon. Our algorithm implements a breadth-first search to check all the possible exon paths from a start codon to an annotated stop codon through a combination of splice junctions. By keeping track of the last junction and the reading frame in a path, LeafCutter2 is able to classify millions of junctions in a computationally efficient manner. LeafCutter2’s algorithm works as follows:

#### Algorithm 1

**junction_classifier**

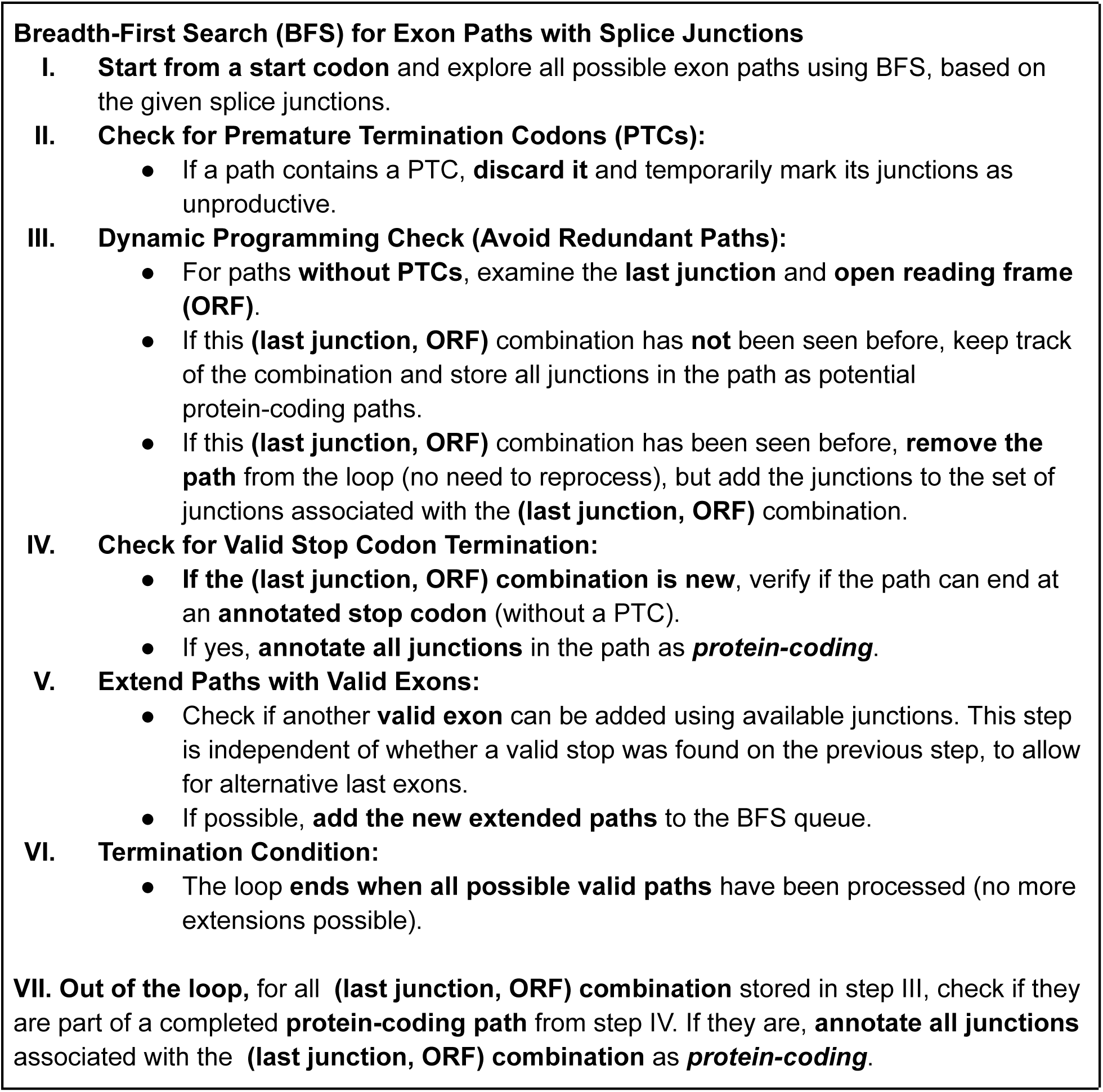
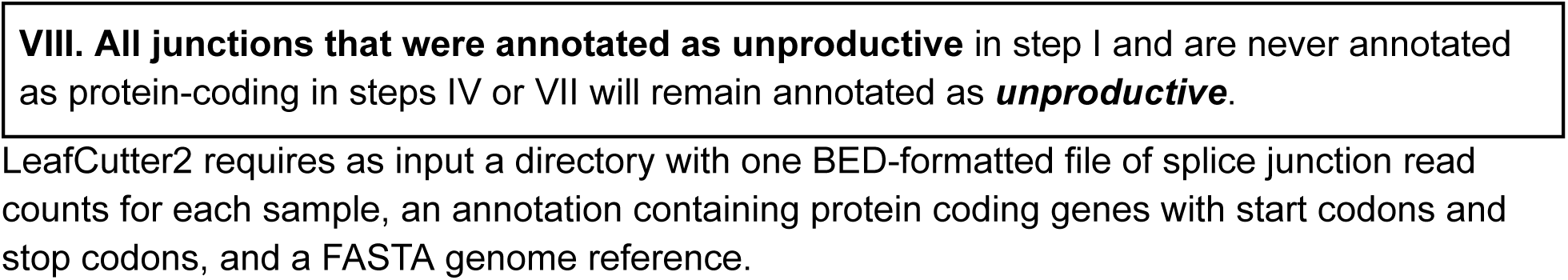

### LeafCutter2 code

A pseudocode of the algorithm is available in **Appendix 1**. LeafCutter2’s junction classifier extension is available for users at https://github.com/cfbuenabadn/leafcutter2.

Two main scripts are implemented in LeafCutter2 for junction clustering and classification:

- `leafcutter_make_clusters.py`: (optional) clusters splice junctions into intron clusters. It helps save time when LeafCutter2 will be run multiple times across different groups.

○ Input:

▪ Junction read counts in BED format for each sample.
○ Output:

▪ `{prefix}/clustering/leafcutter2_pooled:` pool of introns from a list of junction files.
▪ `{prefix}/clustering/leafcutter2_clusters:` intron clusters refined by filters such as minimal junction reads, minimal cluster reads or ratios.
- `leafcutter2.py`: The main LeafCutter2 script, it clusters (unless clusters are provided as input), classifies and quantifies splice junctions into intron clusters labeled as productive, unproductive, or near untranslated region. This script replaces LeafCutter’s `leafcutter_cluster_regtools.py` script.

○ Input:

▪ Junction read counts in BED format for each sample.
▪ GTF annotation with genes, start codons and stop codons.
▪ Genome sequence in FASTA format.
▪ `clustering/leafcutter2_clusters` (optional). The output of `leafcutter_make_clusters.py`. If provided, the script will use these clusters and skip the clustering step.
○ Output:

▪ `leafcutter2.cluster_ratios.gz.` A phenotype matrix with splice junctions, intron clusters, and read counts for each sample divided by the total number of counts per cluster.
▪ `leafcutter2.junction_counts.gz.` A phenotype matrix with splice junctions, intron clusters, and read counts for each sample. This file can be used as input for LeafCutter’s `leafcutter_ds.R` differential splicing script, or to create phenotype tables for QTL calls.
▪ `clustering/leafcutter2_clusters` (optional). Intron clusters. Only created if no intron clusters are provided as input.

junction_classifier algorithm is implemented in LeafCutter2’s `leafcutter2.py` script. The output of this script includes a Boolean value for junction_classifier algorithm. However, junction-level classifications can be further enriched by transcript annotations, which we convey in the output as additional Boolean bitwise flags for each junction. The Boolean bitwise flags reported by the script are, numbered by right-to-left bit-position: (0) UTR (True if junction span is outside range of most downstream START and most upstream STOP) (1) LF2 annotated (True if junction can combine with other observed or annotated productive junctions to create a START to STOP spanning ORF), (3) IsAnnotated (True if junction is in any annotated transcript) (3) IsProductiveAnnotated (True if junction is any annotated productive transcript). For example, the bitflag 1101 (decimal representation 13) corresponds to a junction that is in a UTR region (thus it would likely create a splice junction greater than 50nt downstream from a stop codon) and the junction_classifier algorithm deems it unproductive, though it is in an annotated productive transcript. For some analyses, we deemed it appropriate to handle these junctions differently than other junctions that the junction_classifier algorithm deems unproductive. When relevant, we indicate bit flag sets used to classify junctions as “unproductive”, “productive”, or “UTR” within the relevant methods section.

### Additional NMD rules

In addition to the junction_classifier algorithm, LeafCutter2 computes three key metrics to stratify unproductive splicing junctions (excluding those flagged as UTRs) based on known features influencing NMD efficiency. This add-on requires transcript annotation e.g. from GENCODE. The three rules are: **(1)** the 50 nucleotide rule identifies junctions that introduce premature termination codons (PTCs) within 50 nucleotides of the penultimate exon’s 3’ end or PTCs that are located in the final exon. We report the maximum distance from the PTC to the last exon-exon junction (allowing the enforcement of the 50-nt rule). **(2)** Number of introns upstream or downstream of the PTC. For each unproductive junction, we determine the median number of exons/introns upstream and downstream of the induced PTC when the junction is mapped onto annotated protein-coding transcripts. **(3)** A check for whether the PTC falls within a long exon (>407 nucleotides). For this, we report the modal exon length (most common length) where the PTC is introduced across all compatible transcripts. Additionally, we required junctions to match a 3’ or 5’ splice site in an annotated protein-coding transcript to be tested. Only PTCs induced by substituting the unproductive junction into these transcripts are considered.

To assess the impact of these rules on inducing NMD, we compared the enrichment of unproductive junctions in nascent RNA-seq versus steady-state RNA-seq, stratifying them by the three NMD-related rules: 50-nt rule compliance, exons before/after PTC, and PTC location in long exons (>407 nt). For each rule, we tested whether unproductive junctions satisfying the rule showed stronger depletion in steady-state RNA (indicating NMD activity) compared to those not satisfying it. As a negative control, we also performed equivalent analyses on productive junctions.

### Differential unproductive splicing in GTEx tissues

Exon-exon splice junction read count data from all GTEx tissues was downloaded from the open access dataset from public release GTEx Analysis V8^35^. We used customized scripts to preprocess the splice junction counts into BED files with the splice junction read counts for each sample. This resulted in 17,350 BED files with splice junction read counts.

To ensure that all pairwise comparisons have the same intron clusters and cluster IDs, we pre-clustered the introns using all 17,350 samples, using LeafCutter2’s auxiliary `leafcutter_make_clusters.py script`.

For downstream analyses, including differential splicing and expression analysis and QTL calling, we selected 49 tissues with at least 80 samples each (total of 17,329 samples). For all possible 1,176 tissue pair combinations with the selected 49 tissues, we first classified and quantified the junction read counts using LeafCutter2’s `leafcutter2.py` script using the following inputs:

- All BED splice junction read counts for the two tissues in the pair.
- The pre-clustered introns created with `leafcutter_make_clusters.py`.
- Gencode’s v43 primary assembly annotation in GTF format (for genes, start codons and stop codons).
- Gencode’s GRCh38 primary assembly FASTA file.

We restricted the analysis to 80 samples per tissue for computational efficiency and to prevent bias from differences in sample sizes, and we considered a maximum of 1000 introns per gene. The group files required for LeafCutter’s differential splicing were generated using a custom script, adding the *log* of the percentage of unproductive splice junction reads in each sample (compared to the sample’s total splice junction read counts) as a confounder of the group files.

We tested for differential splicing for each pair of tissues using standard LeafCutter’s `leafcutter_ds.R` script with version 0.2.9, using the count files generated from the previous step. We required each intron to have at least 5 counts in at least 5 samples in total, and at least in three samples per tissue. We corrected for multiple tests using the Benjamini-Hochberg false discovery rate correction.

### Differential gene expression in GTEx tissues

Raw gene read counts from all samples across all GTEx tissues were downloaded from the open access dataset from public release GTEx Analysis V8. We tested for differential expression using between all 1,176 possible pairs of the previously selected 49 tissues using DESeq2^36^ version 1.32.0, and corrected for multiple tests using the Benjamini-Hochberg false discovery rate correction.

TPM data used for visualization of gene expression and for correlation between NMD factor expression and the percentage of unproductive splice junction reads was also downloaded from the GTEx open access datasets. RNA Integrity Number for each sample was obtained from the RNA-seq metadata in GTEx.

### Clustering and gene ontology enrichment analysis

We identified 5,366 significant differentially spliced unproductive clusters at an FDR of 10% and a minimum delta PSI of 0.1. 4644 of these clusters were hosted by genes that were differentially expressed at an FDR of 10% and a minimum log2 fold change of 1. 2418 of these clusters, containing a total of 4103 unproductive splice junctions, were present in all 49 tissues. We used hierarchical clustering from R’s ComplexHeatmap library on the Z-score of the median PSI of each unproductive intron per tissue. We identified six clusters, which were grouped into three super clusters: unproductive splice junctions associated with brain tissues, testis, and other tissues.

Gene ontology enrichment analysis on the genes hosting the unproductive splicing junctions in each cluster was performed with GSEApy version 1.1.4, using MSigDB’s C5 Gene Ontology Biological Process signature set.

### Data processing for HeLa NMD knockdown, LCL chromatin-associated RNA-seq, and FRAC-seq samples

Short-read RNA-seq FASTQ files from HeLa cell samples with shRNA double knockdown of *SMG6* and *SMG7*, as well as shRNA controls, were downloaded from a previous study by Karousis and colleagues^18^ (SRA accession SRP083135). Reads were aligned to Gencode’s GRCh38 primary assembly of the human genome using STAR version 2.7.7a. LeafCutter2’s clustering and classification of splice junctions, as well as differential splicing between *SMG6*/*SMG7* double knockdown and control samples, was performed as in the case of the GTEx samples, except requiring only 2 samples per intron to adjust for the smaller number of samples per group.

The same pipeline was used to process other NMD factor knockdown data from Karousis and colleagues^18^, chromatin-associated RNA-seq from 86 lymphoblastoid cell lines^1^, and 24 FRAC-seq samples^37^. Junctions for all samples were extracted, clustered, and quantified using the Leafcutter2 pipeline. These quantifications were used for all analyses presented in Fig. 1.

### Comparative RNA-seq analysis data processing

RNA-seq reads generated from Cardoso-Moreira et al.^38^ were downloaded from ENA (project accession numbers: PRJEB26695, PRJEB26840, PRJEB26869, PRJEB26889, PRJEB26956, PRJEB26969, PRJEB27035), with exact sample names and accessions for the samples used in this study listed in **Supplementary data file 1**.

Reads were trimmed of adapters using fastp^39^ (v 0.20) with default settings, and aligned to their respective reference genome (**Supplementary data file 2**) with STAR^40^ (v 2.6.1) with the following parameters: {--outFilterType BySJout --outFilterMultimapNmax 20 --alignSJoverhangMin 8 --alignSJDBoverhangMin 1 --outFilterMismatchNmax 999 --outFilterMismatchNoverReadLmax 0.04 --alignIntronMin 20 --alignIntronMax 1000000 --alignMatesGapMax 1000000 --twopassMode Basic}. Splice junction reads were extracted as described for HeLa NMD knockdown data processing, and clustered for each species using LeafCutter^13^ (v. 1) with ‘-p 0.0001’, and the leafcutter2 junction_classifier script was used to generate bitwise flags for each junction (see Leafcutter2 code section above). To run the junction_classifier script with annotations from other sources than Gencode, a gtf file must be reformatted to contain “gene_name”, “gene_type”, “transcript_name”, and “transcript_type” tags, where the gene_type and transcript_type are “protein_coding” for annotated productive transcripts. We used a custom script to reformat gtf files from various sources.

To quantify the total fraction of reads that are productive/unproductive, the junction_classifier script was run using all junctions (including constitutive and extremely rare junctions) rather than just those that are clustered by default, while other analyses consider only clustered junctions. For this analysis, ambiguous UTR junctions (classify_junction script decimal flags in {1,4}) were filtered out, and junctions were considered productive if classify_junction script bit0 or bit3 were True, meaning either the algorithm deems junction as productive and/or the junction is in annotated in productive transcript. Junctions were considered adequately expressed to tally towards “unique junctions” (Fig 3b, Supplementary Figure 8b) if after filtering out UTR junctions, the total junction read count across all samples within the species is >0.02 reads per million junction reads, which roughly corresponds to roughly at least 50 junction reads for most species (though it will vary depending on each species’ total sequencing total depth).

### Differential splicing and expression across tissues and developmental stages within each species

FeatureCounts (v. 1.6) was used to generate gene count tables. Log2FC for splicing and host gene expression was estimated using leafcutter_ds.R and edgeR^41^, respectively for each comparison/contrast. Contrasts included comparing each adult tissue pair to one another within each species, comparing neonate vs adult within each tissue within each species, and comparing early embryo to adult within each tissue within each non-human species. A full list of which samples are utilized for each contrast is in **Supplementary data file 3**. When comparing logFC of splice junctions to logFC of host genes, we only considered the splice junctions that are in a LeafCutter significant cluster (FDR<5%), with deltapsi>5%. Clusters were considered unproductive if they contained at least one significant unproductive junction (unproductive defined as classify_junction decimal flag in {0,1,4,5}). Only one representative splice junction was considered for each cluster, prioritizing the largest absolute deltapsi in each significant cluster.

### Identification of conserved splicing events

We first created a mapping of each splice junction in a query species to the orthologous splice junction (if it exists) in human: In each non-human species, a bed12 file was created for each unique splice junction where each feature (row) contains defines a junction and 1bp flanking on each side, with the 1bp on each side being the feature’s “blocks” that are to be lifted over. CrossMap.py (v 0.6) was used to liftover the bed12 file with UCSC chain files to hg38, to identify the orthologous splice junction in human. Query junctions were discarded if the orthologous splice junction in human was not observed as a spliced junction read. Further filters to identify conserved events consider if the query species junction and its human counterpart are unproductive (classify_junction decimal flag in {0,1,4,5}), if it is alternatively spliced at an appreciable rate (present in a leafcutter cluster), and if it is differentially spliced in human (FDR<5% with abs(deltapsi)>0.05) and the query species (nominal P<0.05, with concordant direction as in the analogous human contrast).

We identified conserved poison cassette exons using a custom script that considers potential pairs of splice junctions corresponding to the two unproductive flanking splice junctions used for exon inclusion. If both flanking junctions are leafcutter2-classified unproductive junctions in humans and one non-macaque species (classify_junction script decimal flags in {0,4,5,8,12}), and the exon-skipping junction is productive (decimal flag in {12,13,14,15}) and they are flanked by annotated coding sequence (as opposed to UTR), and the two junctions are less than 500bp apart, then then the intervening exon is considered a conserved cassette poison exon. As a control, we similarly identified conserved cassette productive exons which are productive when included or excluded in both species (flanking junction decimal flags in {14,15}, and exon-skipping junction decimal flags in {12,13,14,15}). To quantify dN/dS and conservation of cassette exons, and their flanking exons as controls, we utilized 100 way vertebrate PhyloP scores^42^ (track downloaded from UCSC, https://hgdownload.cse.ucsc.edu/goldenpath/hg38/phyloP100way/) and 100 way vertebrate whole-genome alignments (downloaded from UCSC, http://hgdownload.cse.ucsc.edu/goldenPath/hg38/multiz100way/). A custom script was used to calculate dN/dS over the relevant exonic intervals, using the appropriate reading frame from the genome alignments. A table of the identified cassette exons (one unique cassette exon chosen per leafcutter cluster if there are more than one set of potential flanking junctions) and their dN/dS and PhyloP scores is in **Supplementary data file 4**.

### sQTL and eQTL calling in GTEx data

To call sQTLs on the 49 selected GTEx tissues, we ran LeafCutter2’s clustering and classification step using the `leafcutter2.py` script. We ran the script for each tissue independently, using all the splice junction read count BED files for each tissue. This ensures that our phenotype matrices contain only clusters that are relevant for each tissue, while also allowing for tissue-specific clusters that might be filtered out when clustering all tissues together.

We used a modified version of the `prepare_phenotype_table.py` script from Leafcutter to work with LeafCutter2’s output. This script selects intron clusters observed in at least 60% of samples across the tissue. Then we calculate the percent spliced-in (PSI) of each junction, and apply Z-score normalization to each junction across all samples. Finally, we apply rank-normalization to each sample across all junctions to ensure normality in the data. We obtained a covariate matrix for each tissue using principal component analysis (PCA) on this matrix, and selecting the number of top principal components (PCs) that explain more variance than the top PC in a permuted version of the matrix.

Next we used the standard and rank-normalized matrix and the covariate matrix as inputs for QTLTools version 1.3.1 for cis-QTL mapping, using a window of 1 mega-base for each splice junction for both the permutation and nominal pass versions. We only tested for SNPs with a minimum allele frequency (MAF) of 0.05 or greeter. For the permutation pass, we used the Benjamini-Hochberg correction on the adjusted β distribution p-values to obtain the False Discovery Rate.

For eQTL mapping, we first normalized the raw counts for each gene into counts per million (CPM). Next we applied the standard and rank normalization to the log_2_ CPM matrix for each tissue individually. We obtained a covariate matrix and performed nominal pass cis-QTL mapping like described for the splicing phenotypes.

### Colocalization of GTEx molecular QTLs and GWAS traits

GWAS summary statistics from 20 complex phenotypes were downloaded from the GWAS Catalog (**Supplementary Table 3**). All SNP coordinates were standardized to the ChGR38 genome assembly using LiftOver when necessary. For each GWAS trait, we selected SNPs with a p-value threshold of 5 × 10^−8^ for colocalization. For each selected SNP, we considered a window of 1 mega-base. Summary statistics of both sQTLs and eQTLs were obtained with QTLTools nominal pass, selecting only those phenotypes within the GWAS SNP windows that had a permutation pass p-value of 0.01 or lower. We used Hyprcoloc with default settings to colocalize the GWAS signals with the sQTLs and eQTLs of each individual tissue at a time.

### Bulk eQTL in three brain regions

We generated bulk RNA-seq data from postmortem samples across the dorsolateral prefrontal cortex (DLPFC), posterior cingulate cortex (PCC), and head of caudate nucleus (AC) of the ROS/MAP cohort. For library preparation, ≥5μg total RNA from each region underwent poly(A) selection (mRNA-seq, N=911), while 500ng total RNA was processed through rRNA depletion (total RNA-seq, N=1650). Libraries were sequenced at 30 to 50 M paired-end reads per sample with exact depth varied by batches. Following standard QC via FastQC^43^ and adapter trimming via fastp^39^, reads were aligned to the human reference genome GRCh38 using STAR^40^ with WASP correction to reduce reference bias^44^, with further quality control via Picard^45^. Gene-level RNA expression is quantified with RNA-SeQC, removing genes if over 20% samples have TPM expression level of 10% or less. Samples-level RNA quality control was performed following methods outlined by GTEx V8^35^ using three metrics to remove outliers: Relative Log Expression, Mahalanobis distance to hierarchical clustering of samples, and D-statistics quantifying average correlation between pairs of samples. Samples were filtered based on the following thresholds: samples with RLE values in the top 5% (RLEFilterPercent = 0.05), D-statistics in the top 5% (DSFilterPercent = 0.05), or a Mahalanobis distance indicating poor clustering with at least 60% of the top 100 most expressed genes (topk_genes = 100, cluster_percent = 0.6) at a significance level of pvalue_cutoff = 0.05 and cluster_level = 5. Low-expression genes were defined as those with TPM < 0.1 in more than 20% of samples (low_expr_TPM = 0.1, low_expr_TPM_percent = 0.2) and were excluded from the analysis. After quality control, transcript abundance (TPM) matrices were quantile normalized. Technical factors (batch, RNA integrity number, post-mortem interval) and biological covariates (sex, age at death) were included, along with the top 15 genotype principal components to account for population stratification. Additional hidden confounders were inferred using PCA on normalized expression matrices with the number of PCs determined by the Marchenko-Pastur limit^46^. *cis*-eQTL mapping was performed using TensorQTL^47^ across a ±1 Mb window around gene transcription start sites fitting a linear regression model with one variant at a time and incorporating the above covariates. An empirical p-value threshold was first obtained from permutation testing, and genome-wide FDR was then controlled with the q-value method^48^ to define the final significance cutoff. In subsequent integrative analyses (e.g., TWAS, colocalization), all variants were included, regardless of their marginal significance.

### DLPFC single-nucleus RNA-seq eQTL for six major cell types

Single-nucleus RNA-seq was performed on 479 DLPFC samples, each processed individually, as described in our previous work^49^. Briefly, frozen tissue (50-100mg) was homogenized, and nuclei were isolated using a detergent-based lysis and sucrose cushion. After filtering for intact nuclei, 10x Genomics platforms were used for single-nucleus library construction and sequencing, yielding a median of 3,824 nuclei per sample. Doublets were identified using the demuxlet algorithm and DoubletFinder, with thresholds optimized for each library, and further refined using high-resolution clustering. Each nucleus was annotated to cell types using canonical marker genes in a semi-supervised clustering approach implemented in Seurat^50^. We focus on six major cell types with abundant cell proportions to robustly perform eQTL calling: excitatory neurons (Exc), inhibitory neurons (Inh), microglia (Mic), oligodendrocyte precursor cells (OPC), oligodendrocytes (Oli), and astrocytes (Ast). Gene expression counts were summed into pseudo bulk data for each donor-cell pair, quantified as counts per million (CPM), followed by the same normalization, covariates and cis-eQTL mapping procedures previously described for brain bulk datasets.

### DLPFC Histone H3K9ac ChIP-seq and haQTL

H3K9ac ChIP-seq was performed on 669 DLPFC samples, using a well-validated anti-H3K9ac antibody (Millipore #06-942). Approximately 50 mg of gray matter tissue was cross-linked (1% formaldehyde) and sonicated prior to overnight immunoprecipitation in preparation for the ChIP assay. Purified DNA was used to construct libraries (including end repair, adapter ligation, and size selection) and single-end 36bp reads were generated. Reads were aligned to GRCh38, and broad peaks were called with MACS2 (broad region q-value < 0.05). We applied stringent per-sample QC requiring ≥15 million unique reads, non-redundant fraction ≥0.3, and fraction of reads in peaks ≥0.05. A union set of peaks (92,401) was formed across individuals for per sample read quantification, which ensures reliability and power by leveraging information across samples. Peaks are normalized using a standard voom-limma^51^ pipeline with batch correction using ComBat^52^. For cis-haQTL mapping, sliding windows were defined based on topologically associating domains (TADs) and their boundaries derived from combined brain and blood Hi-C data^53^, resulting in 1,328 sliding autosomal TADs with median size of 2,076 kb (range: 1,160-17,363 kb).

### AD GWAS and QTL integration methods

We constructed a custom LD reference panel from the Alzheimer’s Disease Sequencing Project (ADSP), consisting of approximately 17,000 whole-genome sequenced (WGS) individuals of European ancestry. Quality control on a recent AD GWAS meta-analysis summary statistics^25^ was then performed against this reference panel, including allele harmonization and LD mismatch detection using SLALOM^54^, with flagged suspicious variants removed. To prioritize causal genes underlying AD associations using sQTL data, we evaluated evidence from TWAS with colocalization at gene level. Specifically we applied PTWAS^26^, which incorporates probabilistic fine-mapped u-sQTLs, including all u-sQTLs within a given gene, to predict AS-NMD, followed by event-trait association testing using standard TWAS statistics based on GWAS summary data^55^. We then evaluate whether AS-NMD and GWAS share putative causal variants using SuSiE-COLOC (COLOC version 5)^27^ to detect potentially multiple independent colocalization signals. Significant colocalization was defined by PP.H₄ > 0.8, and colocalization sets were determined by accumulating variants ranked by PIP until the cumulative PIP exceeded 0.95 (similar to the 95% colocalization confidence sets in ColocBoost^29^). Gene-level evidence from these analyses was consolidated using significant signals from both colocalization and PTWAS to assess causality for each gene, with those exceeding 28 considered high-confident risk genes for downsteam follow-up^26^.

To integrate molecular QTL data across multiple contexts along with u-sQTL, we applied ColocBoost^29^, a multi-task learning method designed to efficiently handle high-dimensional, multi-modal colocalization analysis while accommodating multiple causal variants per locus. Our ColocBoost analysis included 16 molecular QTL datasets, comprising bulk eQTLs from the DLPFC, PCC, and AC, u-sQTL and p-sQTL QTLs from the same regions, DLPFC pQTL, and six pseudobulk eQTLs from distinct DLPFC cell types previously described, along with AD GWAS summary statistics. The results were summarized as 95% colocalization confidence sets (CoS) containing a number of putative causal variants in high LD, which collectively define the most probable shared causal variants for each molecular trait and disease association signal.

### Mediation analysis

Mediation analysis was conducted to investigate the potential indirect effect of the unproductive sQTL on the gene expression level of targets through a splicing event. Residuals of gene expression and splicing levels, corrected for known covariates, were derived from previous eQTL and sQTL analyses. Using the mediation package in R^56^, the analysis was performed in both directions to assess whether there is a true causal relationship.

Specifically, we tested (1) the splicing event as a mediator between the genetic variant and gene expression, and (2) gene expression as a mediator between the genetic variant and the splicing event. For each direction, a two-step regression approach was applied: a generalized linear model (GLM) was first fitted to model the mediator as a function of the genetic variant, followed by a GLM to model the outcome as a function of the genetic variant and the mediator. Covariates, including principal components derived from genotype and molecular data, as well as other factors such as age, sex, and PMI, were included in both models. Bootstrap sampling with 1,000 iterations was used to estimate the indirect, direct, and total effects with robust standard errors. A lack of evidence for causal mediation is defined as the absence of statistically significant indirect effects in either direction of the tested model.

## Code availability

Pipelines and all original code are available on GitHub at https://github.com/cfbuenabadn/leafcutter2/.

## Acknowledgements

We thank members of the Li and Wang labs for discussions and support. We thank N. Gonzales and F. Naumann for their careful reading of our manuscript and their insightful comments. We thank the members of the Alzheimer’s Disease Sequencing Project Functional Genomics Consortium. This work was supported by National Institute of Health grants R35 GM153249 (Y.I.L., B.F., and J.L.), R01 HG011067 (Y.I.L. and B.F.), R01 AG076901 (G.W. and X.C.), a FunGen-AD Consortium Grant U01 AG072572 (G.W., F.R. and P.L.D.), a GREGoR Consortium Grant (Y.I.L. and C.F.B.A.N.), and R35 GM13808 (X.L. and J.L.).

## Contributions

C.F.B.A.N., R.F., C.D. contributed equally. Y.I.L., G.W. jointly supervised research. C.F.B.A.N., B.F., G.W., Y.I.L.. conceived and designed the experiments. C.F.B.A.N., R.F., C.D., B.F., Q.H., Y.l.L. performed analyses and contributed code to LeafCutter2. J.L. and X.L. contributed statistical methods. C.F.B.A.N., R.F., B.F., G.W. and Y.I.L. wrote the paper.

## Supplementary figures

**Supplementary Figure 1.**
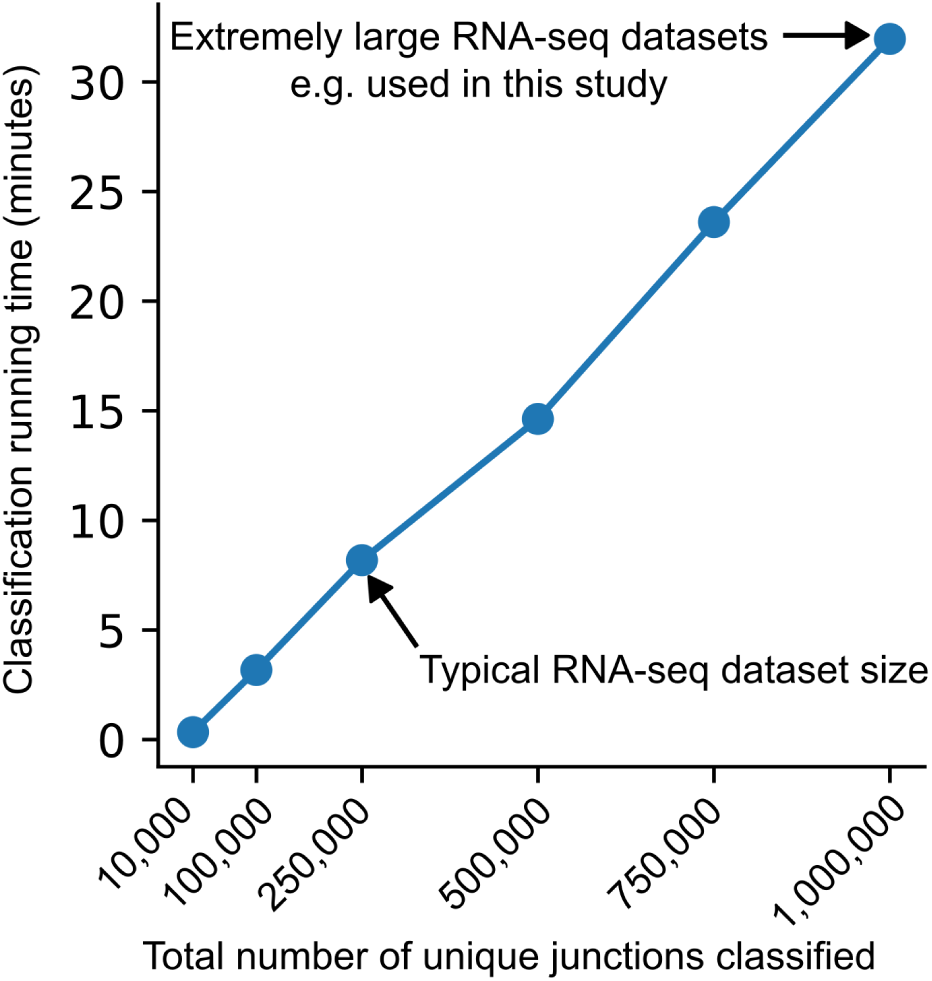
LeafCutter2’s junction classifier runtime as a function of the total number of splice junctions. Most datasets are expected to take approximately 10 minutes longer to be processed compared to the original LeafCutter pipeline. Large datasets with one million splice junctions will take 30 minutes longer.

**Supplementary Figure 2.**
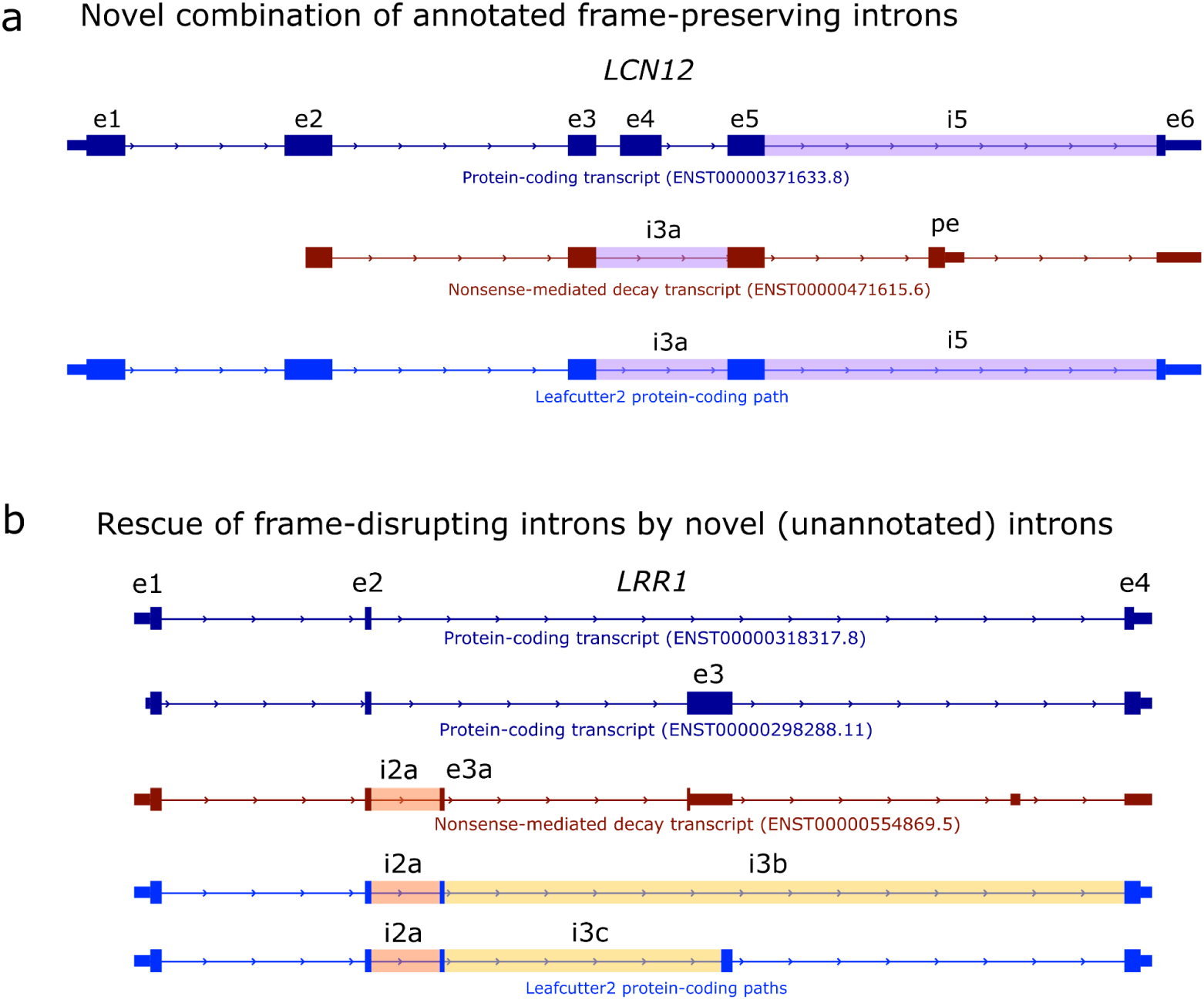
Examples of splice junctions exclusively annotated in Gencode NMD transcripts, but classified as productive by LeafCutter2. **a)** The splice junction (i3a) skipping exon 4 (e4) in *LCN12* is only annotated in the transcript ENST0000047165, which is annotated as a nonsense-mediated decay transcript. However, the premature termination codon in this transcript comes from a downstream poison exon (pe). Since e4 is a symmetric exon (its length is a multiple of 3), i3a does not disrupt the reading frame, which allows LeafCutter2 to find a novel protein-coding path between the start codon in exon 1 (e1) and the stop codon in exon 6 (e6). **b)** The introduction of an alternative exon (e3a) between protein coding exons 2 (e2) and 3 (e3) in *LRR1* disrupts the open reading frame. This creates a premature termination codon in e3, creating a premature termination codon, which targets transcript ENST00000554869 for nonsense-mediated decay. As a result, the splice junction i2a is annotated as unproductive in Gencode. However, two splice junctions (i3b and i3c) missing from the annotation but present in nascent RNA LCL data^1^ can rescue the reading frame disruption caused by i2a, creating a protein coding path between the start codon in exon 1 (e1) and the stop codon in exon 4 (e4)

**Supplementary Figure 3.**
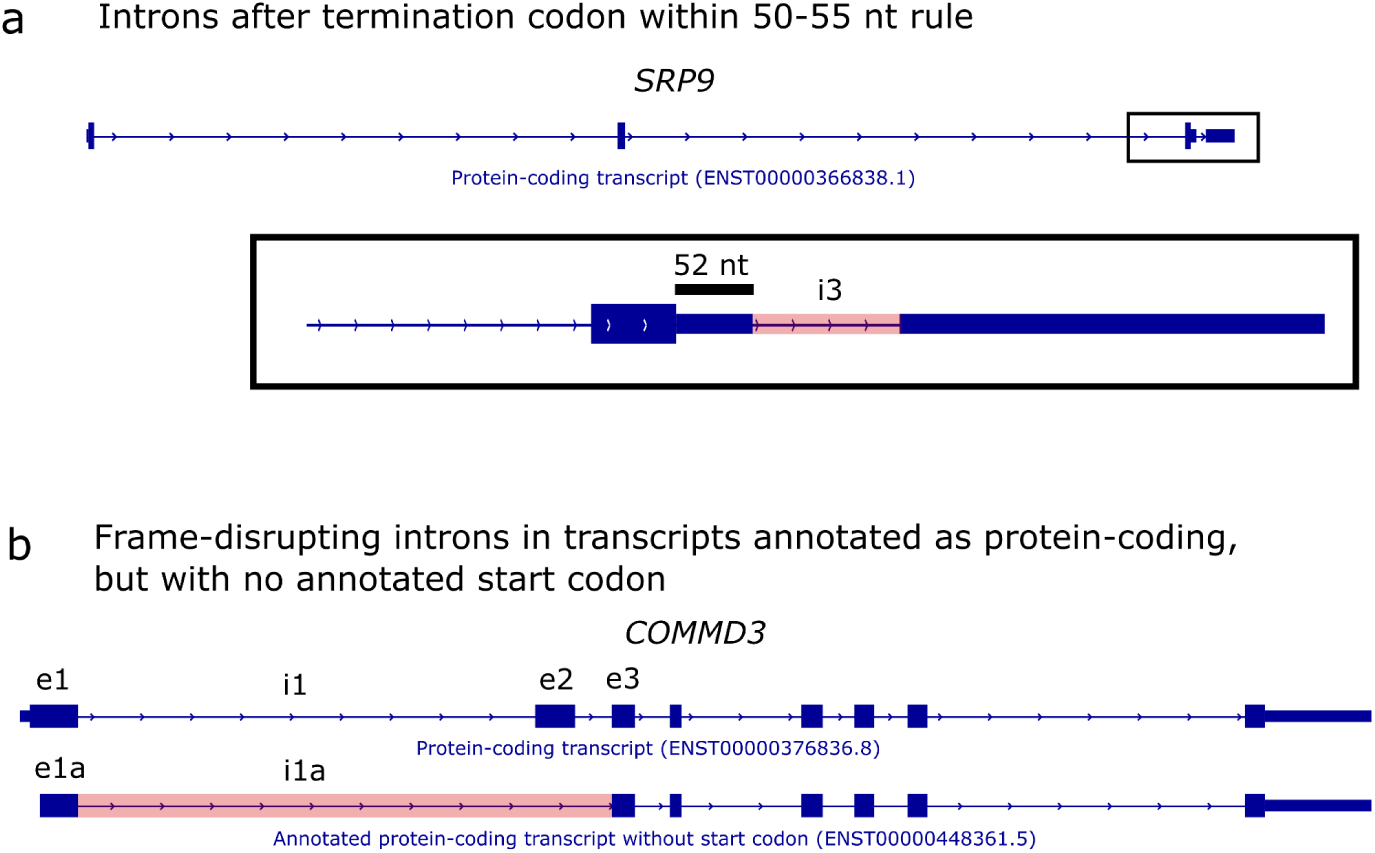
Examples of splice junctions in Gencode protein coding transcripts that are classified as unproductive by LeafCutter2. **a)** Splice junction i3 in *STP9* falls after the stop codon on the last coding exon, but within the 50-55 nucleotide rule. LeafCutter2 misclassifies this splice junction as unproductive. **b)** Transcript ENST00000448361 in *COMMD3* is annotated as protein-coding, but its Gencode annotation lacks a start codon. Intron i1a in this transcript skips exon 2 (e2), an asymmetric exon (length is not a multiple of 3) in other protein-coding transcripts with annotated start and stop codons. Because of the missing start codon and the reading frame disruption, LeafCutter2 is unable to find a viable protein-coding path that includes splice junction i1a.

**Supplementary Figure 4.**
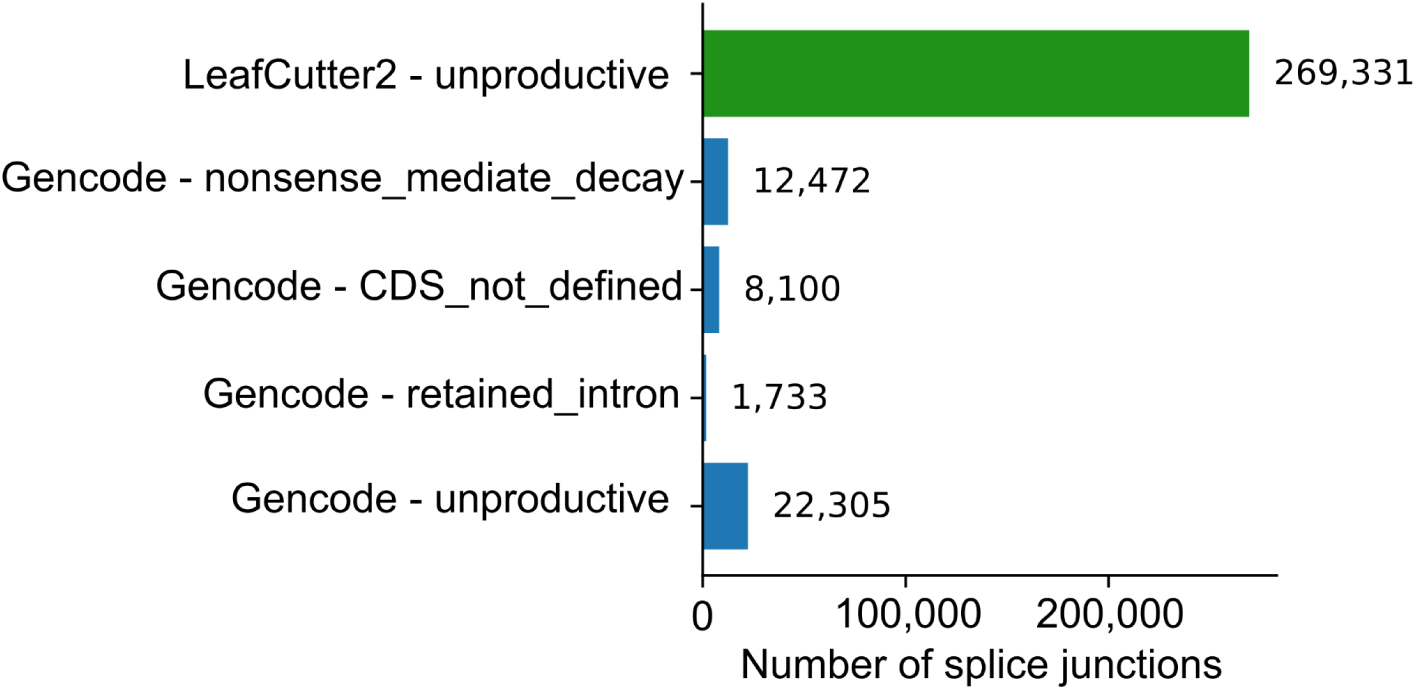
LeafCutter2 finds hundreds of thousands novel unproductive splice junctions in our dataset of 86 nascent RNA samples across 86 Lymphoblastoid cell lines. Most of these are novel. In contrast, Gencode only has 22,305 unproductive splice junctions across multiple categories.

**Supplementary Figure 5.**
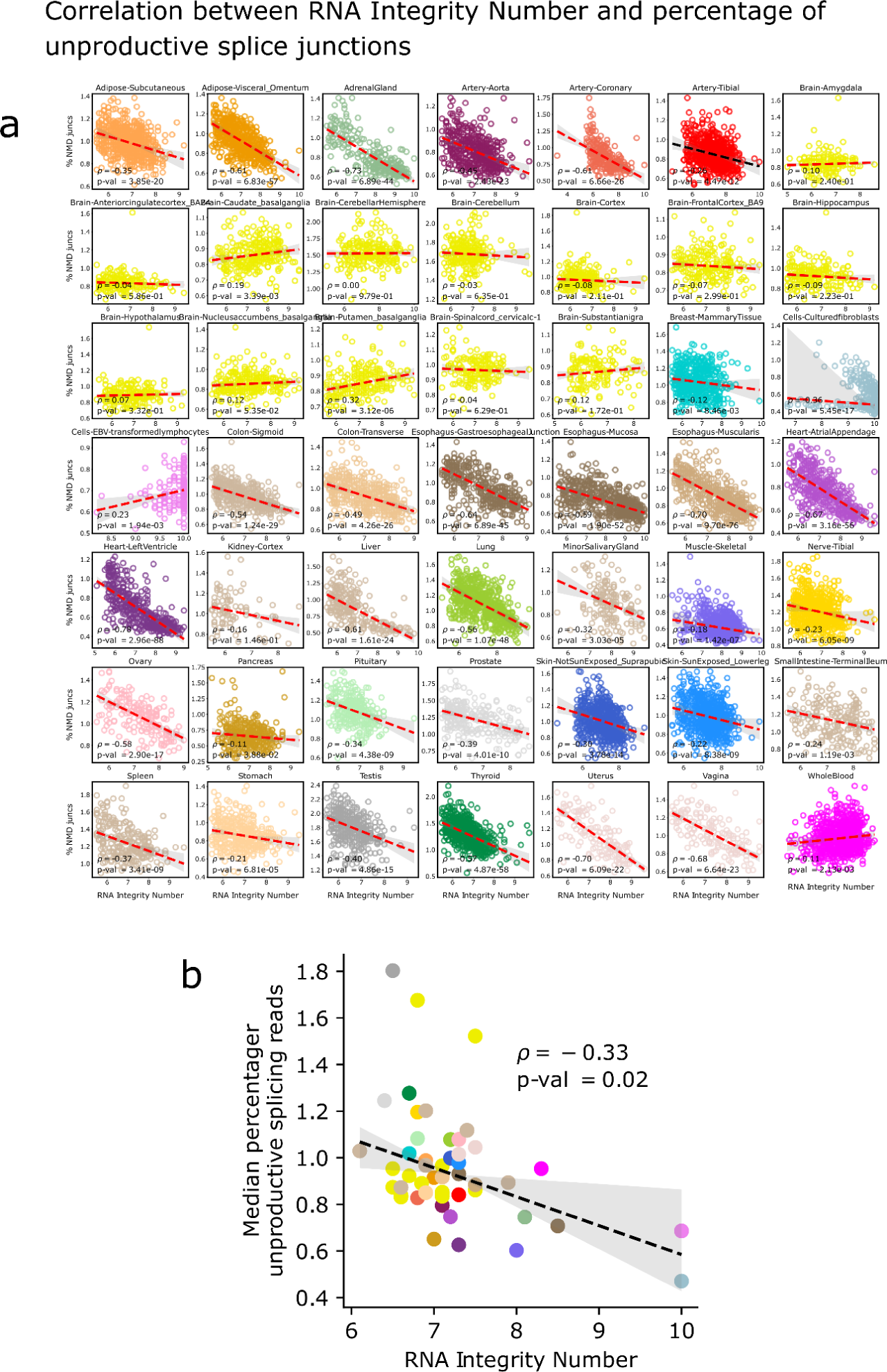
Correlation between RNA Integrity Number (RIN) and percentage of unproductive splice junctions. **a)** Per sample within the same GTEx tissue, and **b)** median across different tissues.

**Supplementary Figure 6.**
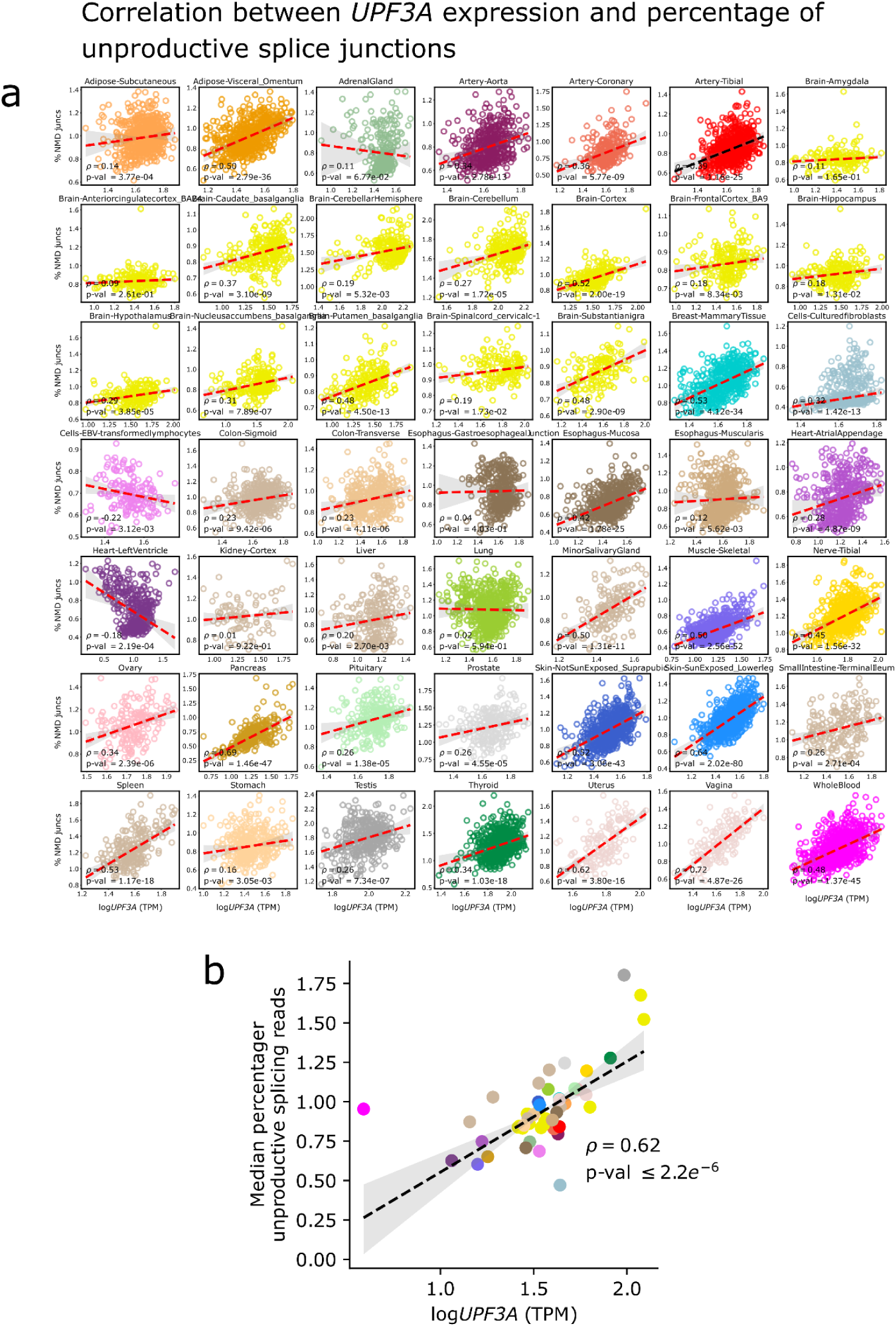
Correlation between *UPF3A* expression and percentage of unproductive splice junctions. **a)** Per sample within the same GTEx tissue, and **b)** median across different tissues.

**Supplementary Figure 7.**
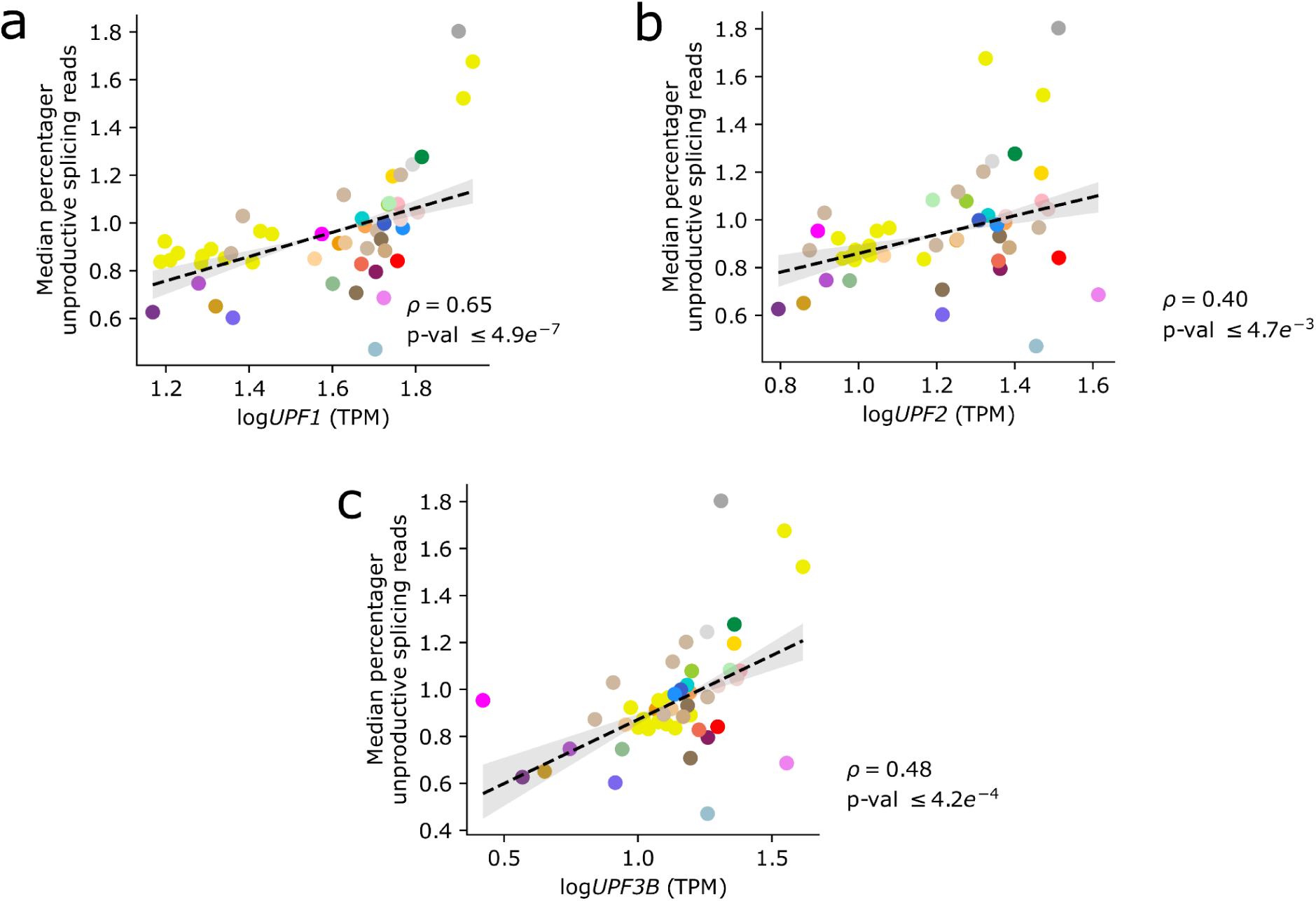
Correlation between NMD factors expression and percentage of unproductive splice junctions. **a)** *UPF1*, **b)** *UPF2*, **c)** *UPF3B*.

**Supplementary Figure 8.**
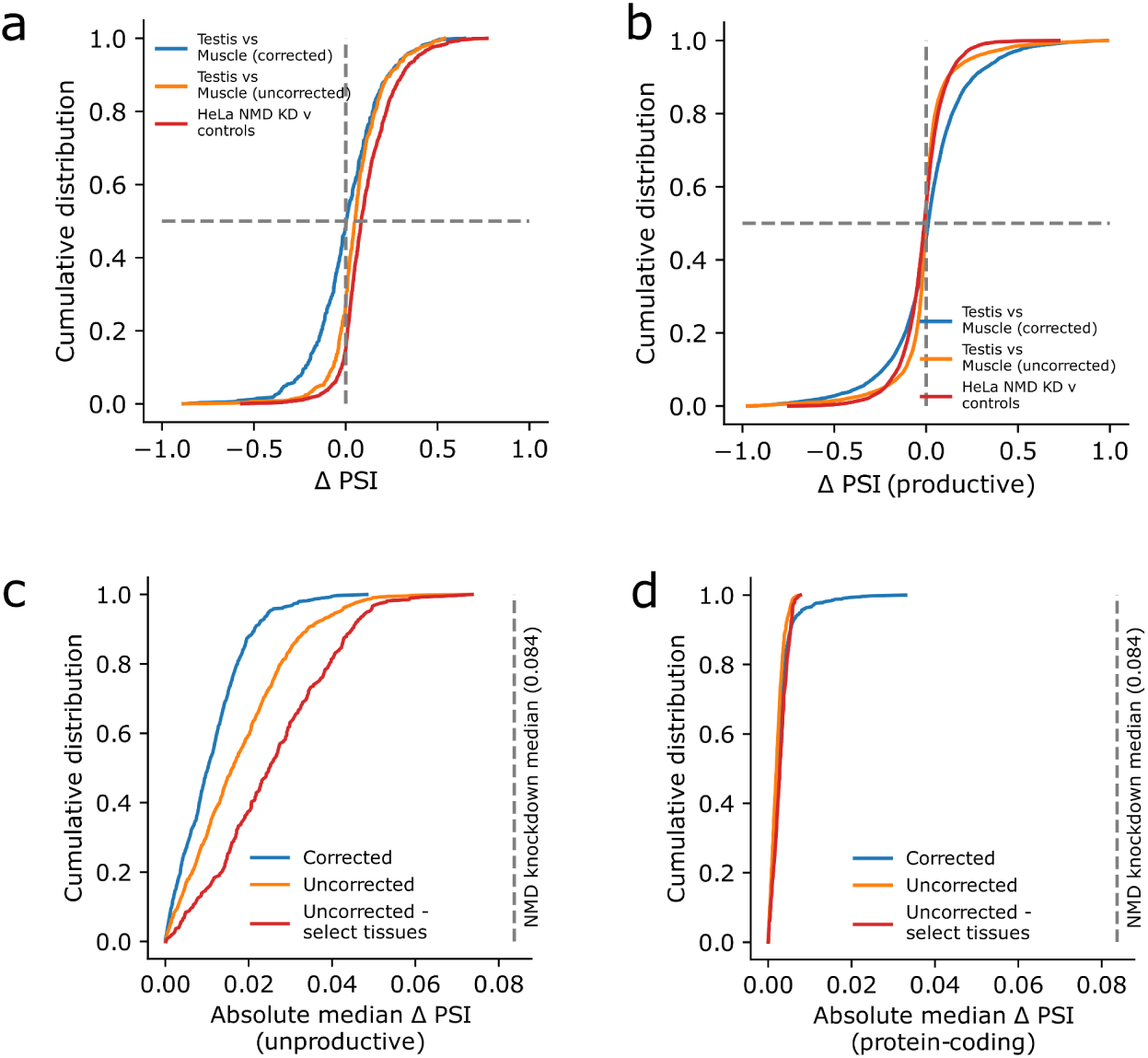
Effect of NMD efficiency and covariate correction in LeafCutter’s differential splicing test. **(a)** Distribution of delta PSI of significant unproductive splice junctions between testis and skeletal muscle when correcting for the percentage of unproductive splice junctions (blue), without correction (orange), and between HeLa cell lines with NMD factors *SMG6/SMG7* double shRNA knockdown (dKD) and controls (red). The dKD versus control comparison serves as an example of contrast that reflects only differences in NMD activity rather than cell-type-specific regulation of unproductive splicing. These data indicate that correcting for the percentage of unproductive splice junctions is required and sufficient to measure unbiased cell-type-specific regulation of unproductive splicing. **(b)** Distribution of delta PSI for productive splice junctions between the same groups. **(c)** Absolute deviation from 0 of the median of the delta PSI distribution of unproductive splice junctions across all tissue pairs when correcting for the percentage of unproductive splice junctions (blue), without correction (orange), and uncorrected when the pairwise comparison contains one of the top three (Testis, Brian - Cerebellum, Brain - Cerebellar Hemisphere) or bottom three (Heart - Left Ventricle, Muscle - Skeletal, Cells - Cultured Fibroblasts) tissues regarding their percentage of unproductive splice junctions (red). **(d)** Absolute deviation from 0 of the delta PSI distribution of productive splice junctions across the same groups. These observations again suggest that differences in NMD activity bias differential splicing measurements of unproductive splicing events, but can be corrected using *UPF3A* expression level as covariate.

**Supplementary Figure 9.**
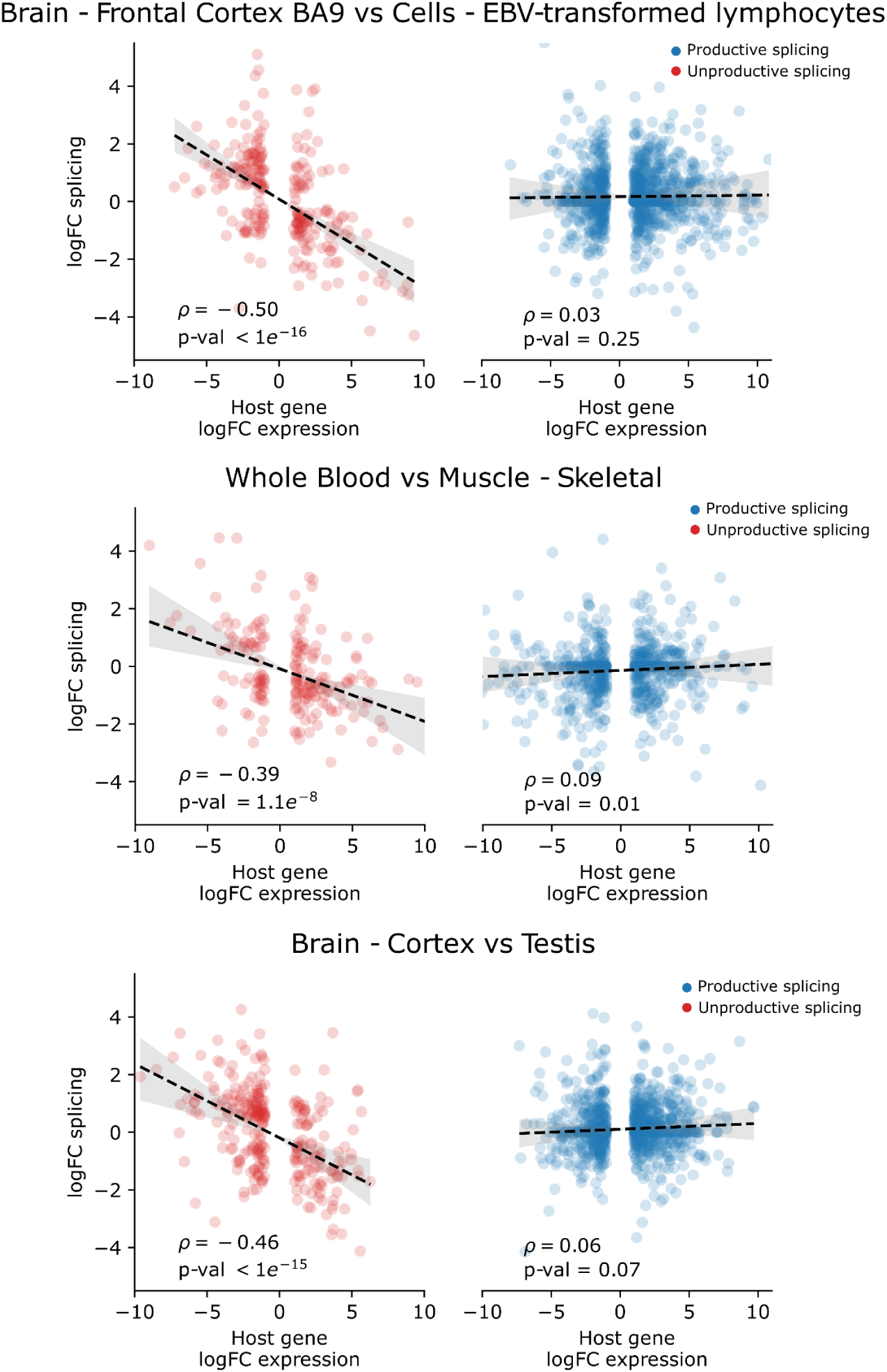
Examples of the negative correlation differential unproductive splicing and differential gene expression across tissue pairs in GTEx. Differential productive splicing has little to no correlation with differential gene expression.

**Supplementary Figure 10.**
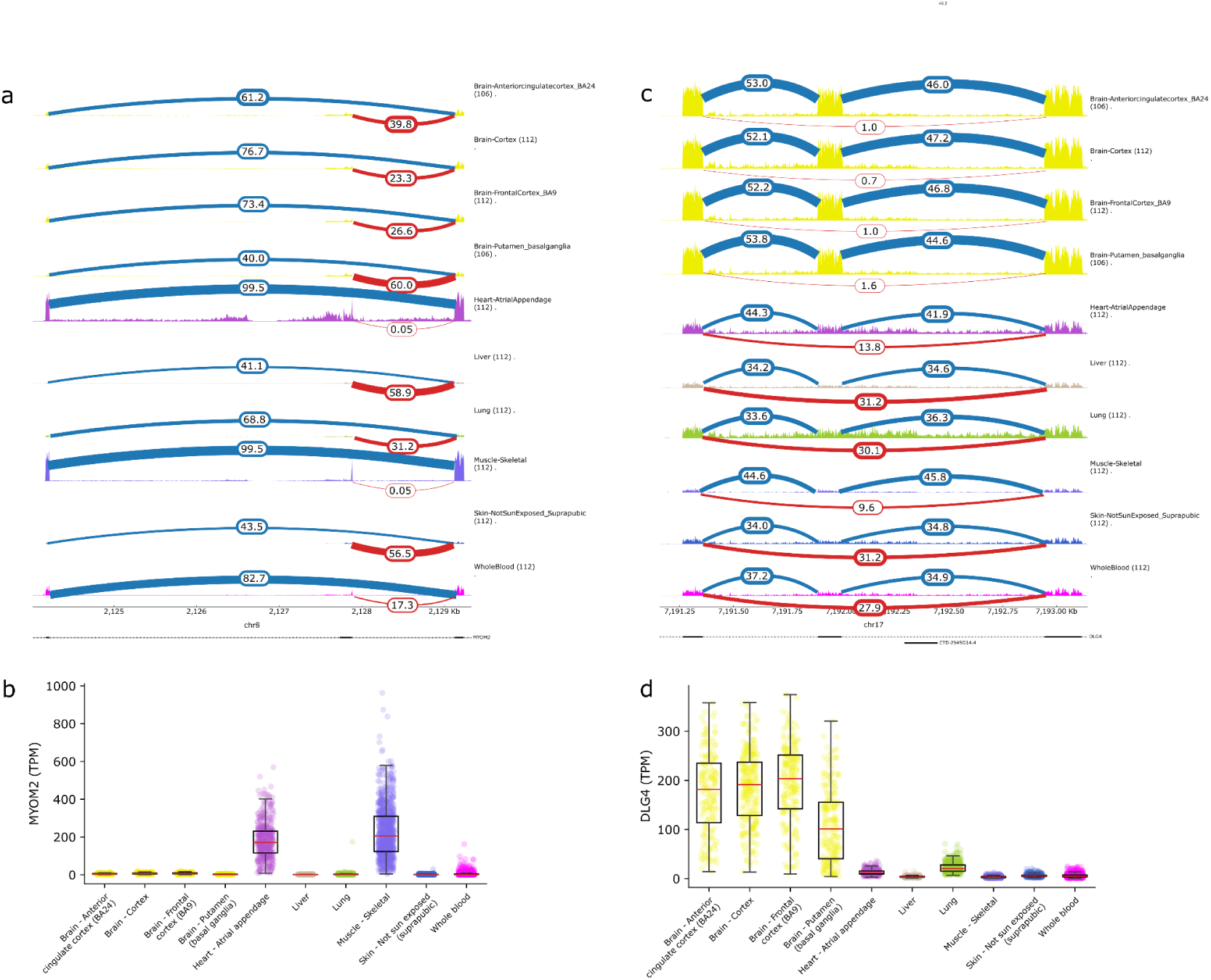
Examples of differential unproductive splicing and its association with gene expression. **a)** Unproductive splicing in *MYOM2* is relatively high in most tissues except skeletal muscle and heart tissue. **b)** *MYOM2* is primarily expressed in skeletal muscle and heart tissues. **c)** Unproductive splicing in *DLG4* is low in brain tissues, but relatively elevated in all other tissues. **d)** *DLG4* is primarily expressed in brain tissues.

**Supplementary Figure 11.**
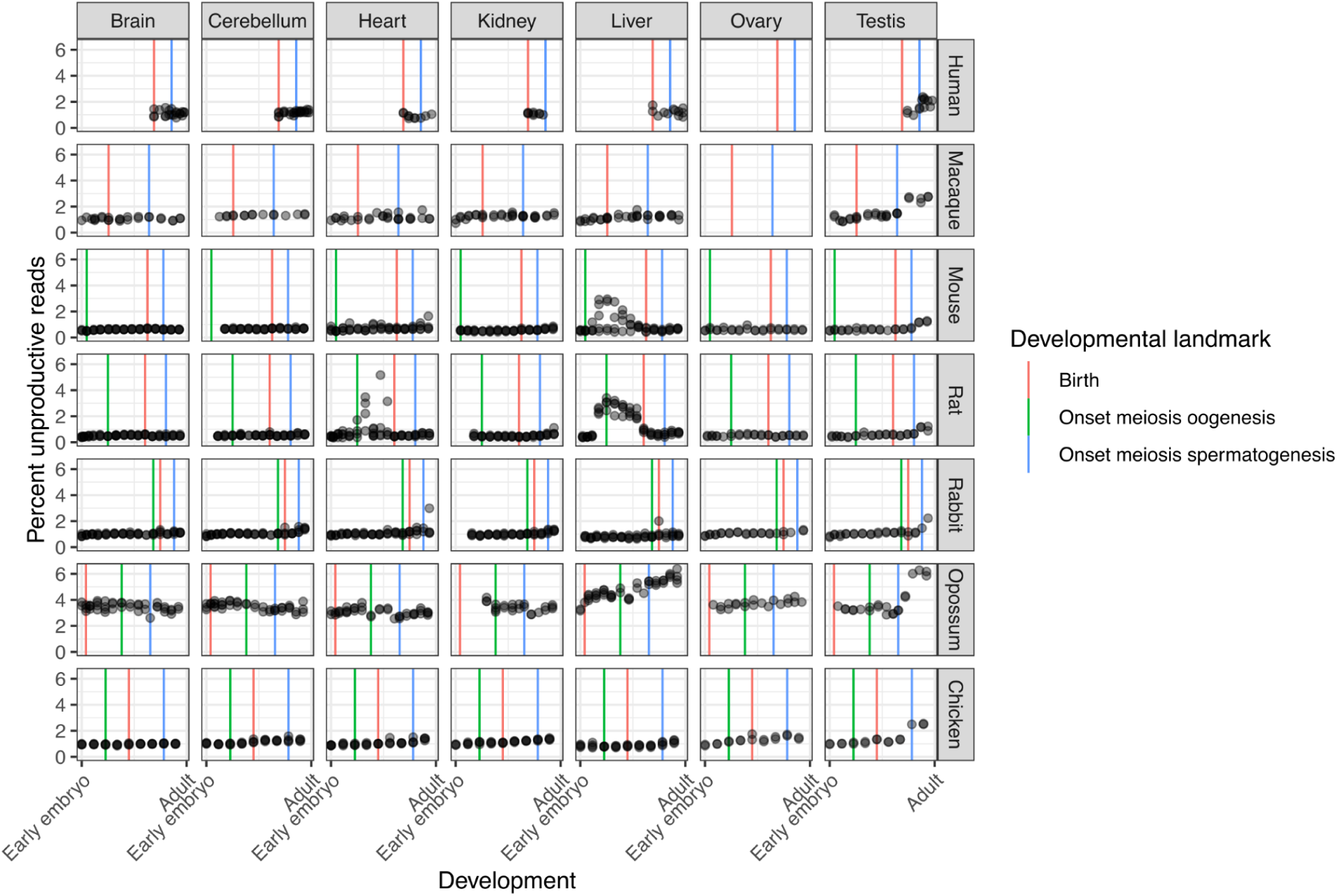
Percent of unproductive reads across species, tissues, and developmental time. RNA-seq samples (points) are ordered by developmental stage within each tissue and species, and the percentage of splice junction reads which are classified as unproductive is plotted. Because samples across species do not necessarily reflect equivalent developmental time courses, we also included developmental landmarks for each of the developmental time courses in each species, as defined by the source publication^38^.

**Supplementary Figure 12.**
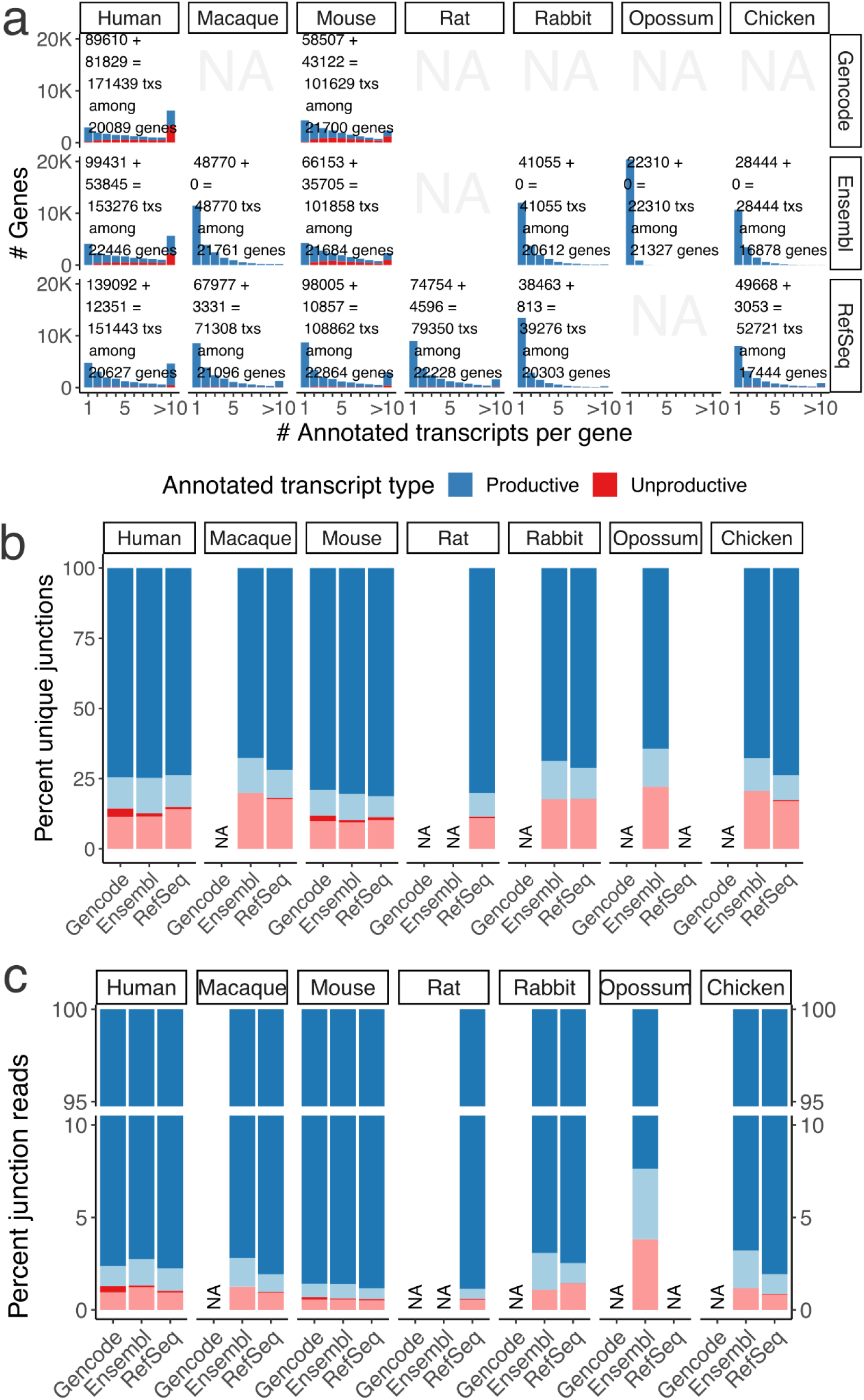
Various sources of gene annotations yield similar fractions of unproductive splice junctions and transcripts. **a**) Histograms quantify the number of transcripts per protein-coding gene in each species’ current gene annotation. The fraction of transcripts that are productive are filled as a fraction within each bin. For each species, gene annotations can be sourced from one or more sources (ie, Gencode, Ensembl, or RefSeq. Some gene annotations not available (NA) from either Gencode or UCSC (which itself sources annotations from Ensembl and/or RefSeq). **b**) Percent of unique junctions (with a minimum abundance of 0.05 junction reads per million junction reads) that are productive or unproductive, and, annotated or unannotated across gene annotation sources. All samples (all tissues and all post-birth developmental time points) are combined for simplicity. **c**) Similar to B, but plotting the percent of splice junction *reads* in each category

**Supplementary Figure 13.**
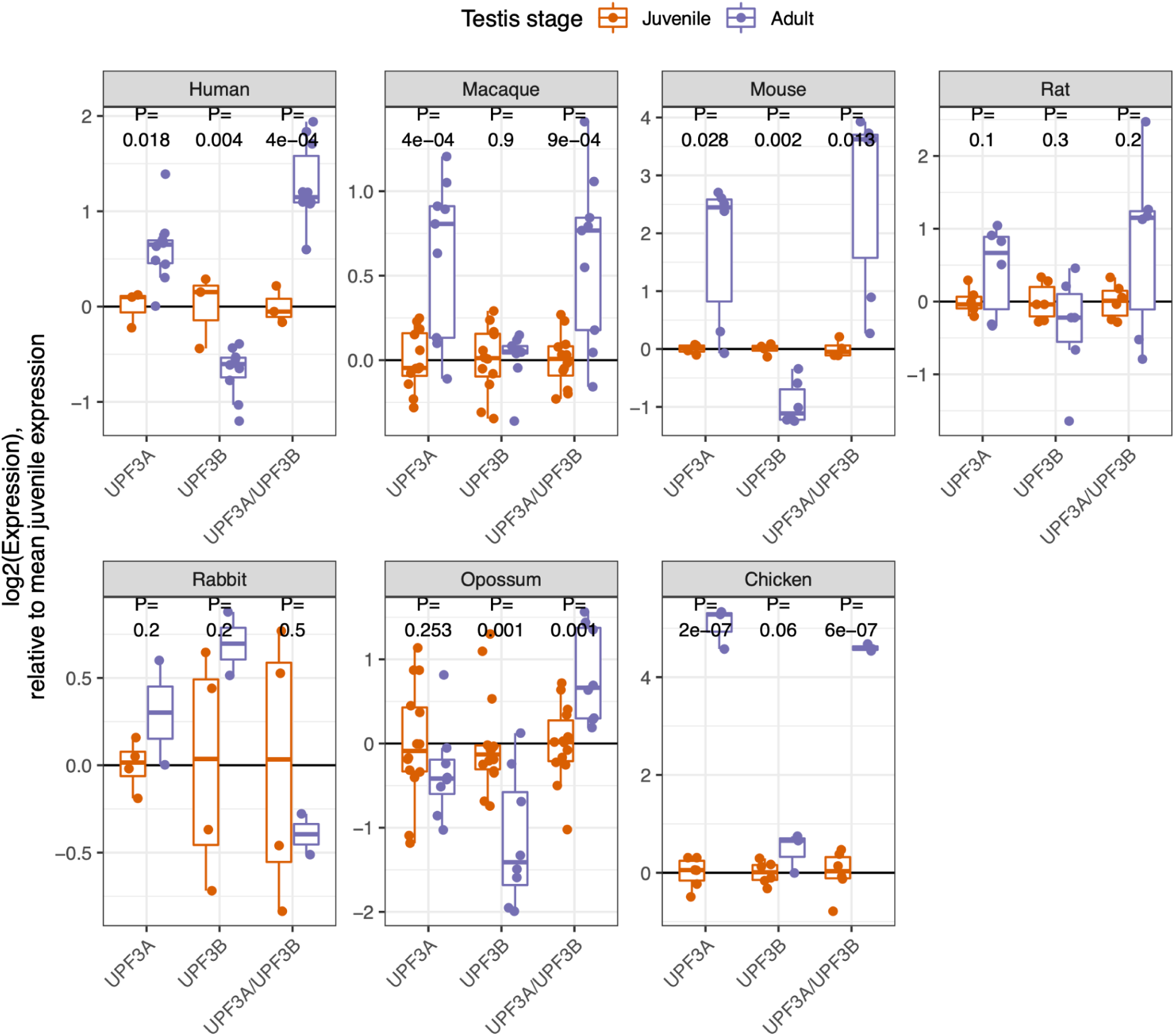
*UPF3A* and *UPF3B* expression across testis development. Boxplots (inner quartiles with whiskers that extend to the most extreme value within 1.5 times the interquartile range from the hinge) and points for individual samples depict the relative expression (RNA-seq) of *UP3A*, *UPF3B*, and the *UPF3A*/*UPF3B* ratio in juvenile and adult testis in each species. P-values from two-sided t-test.

**Supplementary Figure 14.**
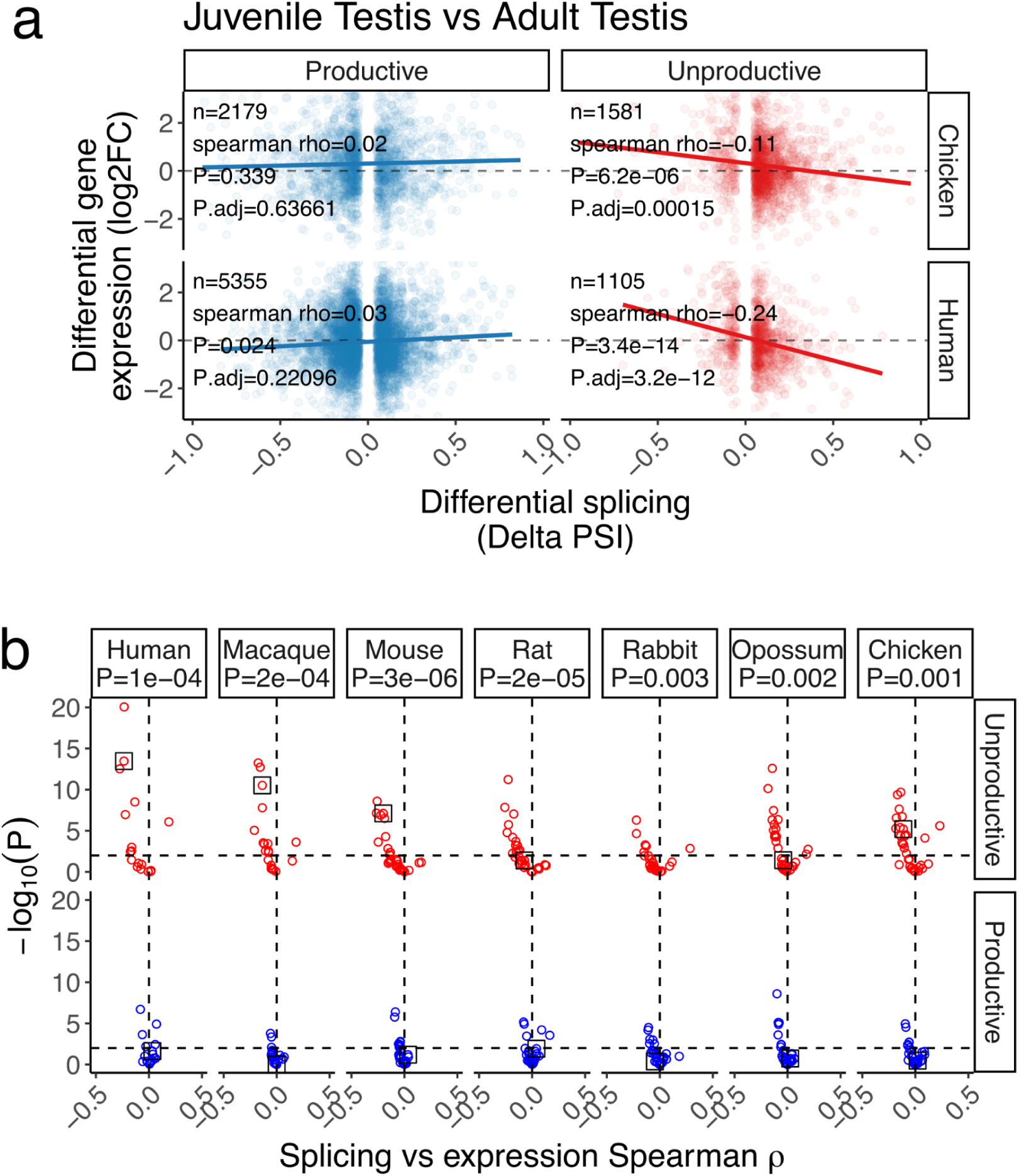
Correlation between splicing and expression for unproductive and productive differentially spliced junctions. **a)** Scatter plots showing the Spearman’s correlation between splicing (delta PSI) and differential gene expression (logFoldChange) for significant differential splicing events (FDR<0.05% and delta PSI > 5%) between juvenile and adult testis in human, grouped by whether the splice event is productive or unproductive. A similar contrast is shown for chicken. P values for spearman correlation coefficient, and adjusted for multiple test correction. **b)** for all contrasts (including comparisons between adult tissue type within each species, and different developemental timepoints within each tissue within each species, see Methods) and the significance and correlation coefficient of the splicing vs expression is shown. Only well powered contrasts with at least 100 significant differentially spliced intron clusters were included. The contrast for juvenile vs adult testis for each species is highlighted with a black box around the point. P-values correspond to a one-sided Mann-Whitney test, comparing the distribution of spearman coefficients for contrasts among productive vs unproductive junctions.

**Supplementary Figure 15.**
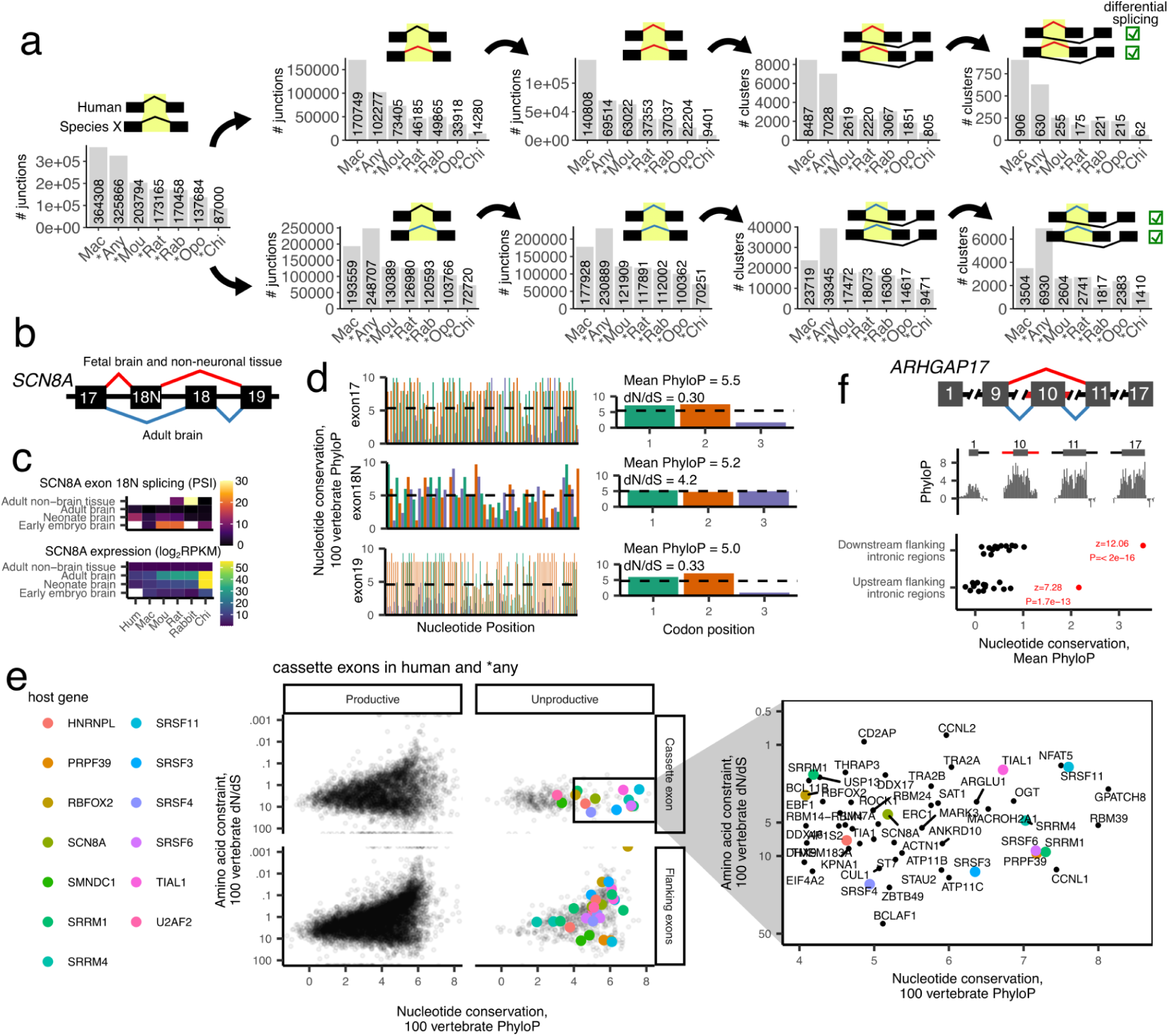
Identification of evolutionarily conserved unproductive splicing events. **a**) Splice events classified as conserved between human and each query species by series of filters. Number splice events (splice junctions, or unique clusters containing at least one splice junction) passing each filter in query species depicted as bars. *Any refers to any non-macaque query species. Filters are: 1) Query-species splice junction is mapped with liftover to splice junction observed in human, 2) splice junction is unproductive in query species, 3) and unproductive in human, 4) and is alternatively spliced in both, 5) in a tissue- or developmental-specific fashion as assessed by leafcutter differential splicing (Methods for details). Filters 2-5 similarly applied for productive splice events (lower row). **b**) Alternative splicing gene-structure of poison exon in SCNA8. **c**) Mean expression and splicing of SCN8A poison exon across samples from various tissues and brain development stages. **d**) Nucleotide conservation (100 vertebrate PhyloP score) across the *SCN8A* poison exon and flanking exons, colored by codon position. Average PhyloP by codon position, and amino acid constraint (dN/dS) plotted on the right for each exon. **e**) Nucleotide conservation and amino acid constraint for unproductive (poison) and productive exons present in human and *any query species (passes filter 4). Each point is an exon. Unproductive cassette exons in genes of interest colored, or labelled in zoomed inset to highlight poison exons with high nucleotide conservation but low amino acid conservation. **f**) Gene structure (top) for unproductive exon skipping junction in ARHGAP17. Nucleotide conservation for 100bp intronic splice site regions (black) flanking each of the ARHGAP17’s 17 exons, highlighting intronic regions flanking the skipped exon (red). P-value represents a two-sided z-test.

**Supplementary Figure 16.**
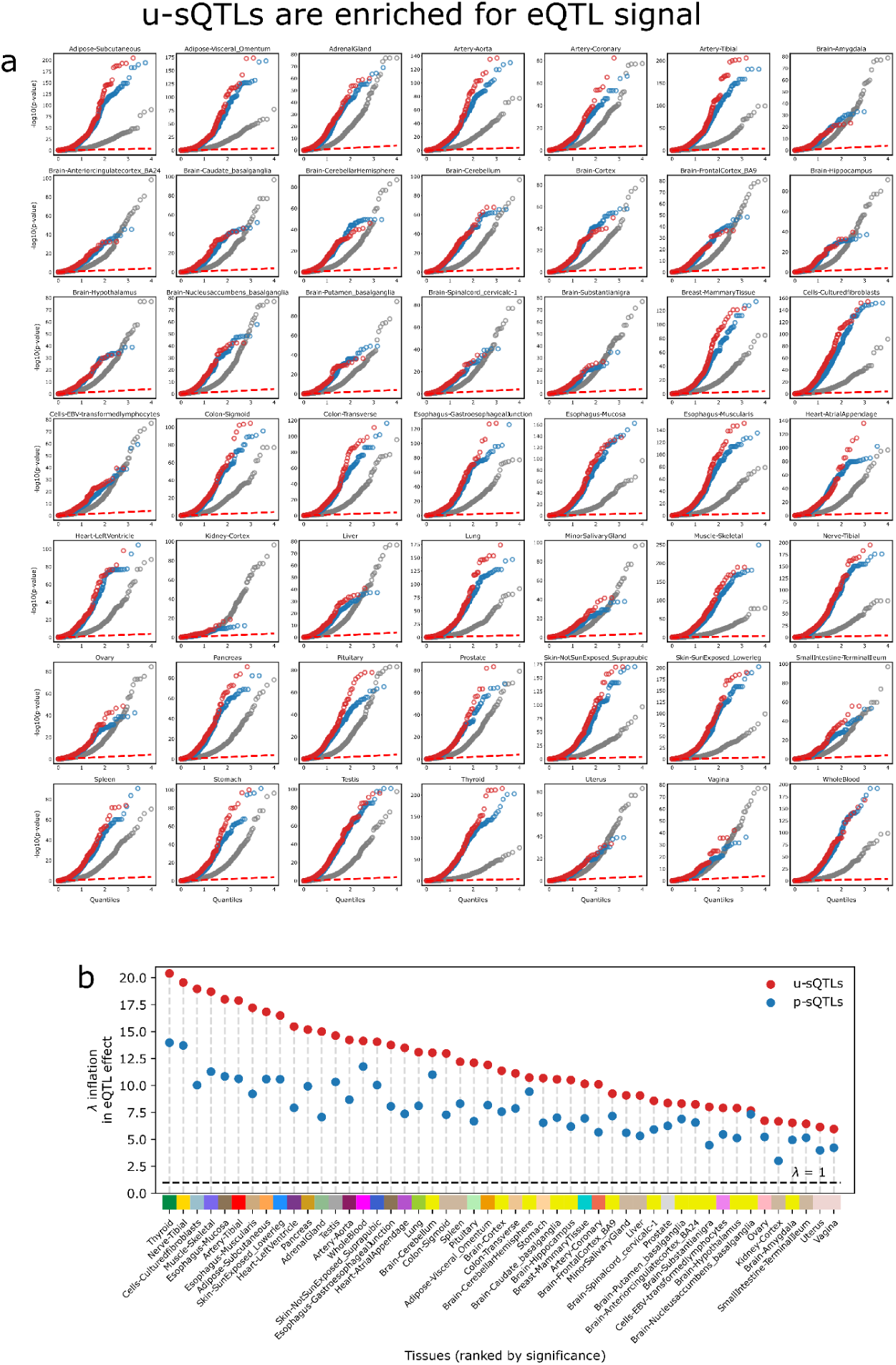
Enrichment of eQTL signals in u-sQTLs. **a)** QQ plots of the eQTL effect across 49 GTEx tissues of u-sQTLs (red), p-sQTLs (blue) and random SNPs (gray). **b)** Lambda inflation shows enrichment in the eQTL p-value distributions of sQTLs, with u-sQTLs signals showing more strength than p-sQTLs across all GTEx tissues.

**Supplementary Figure 17.**
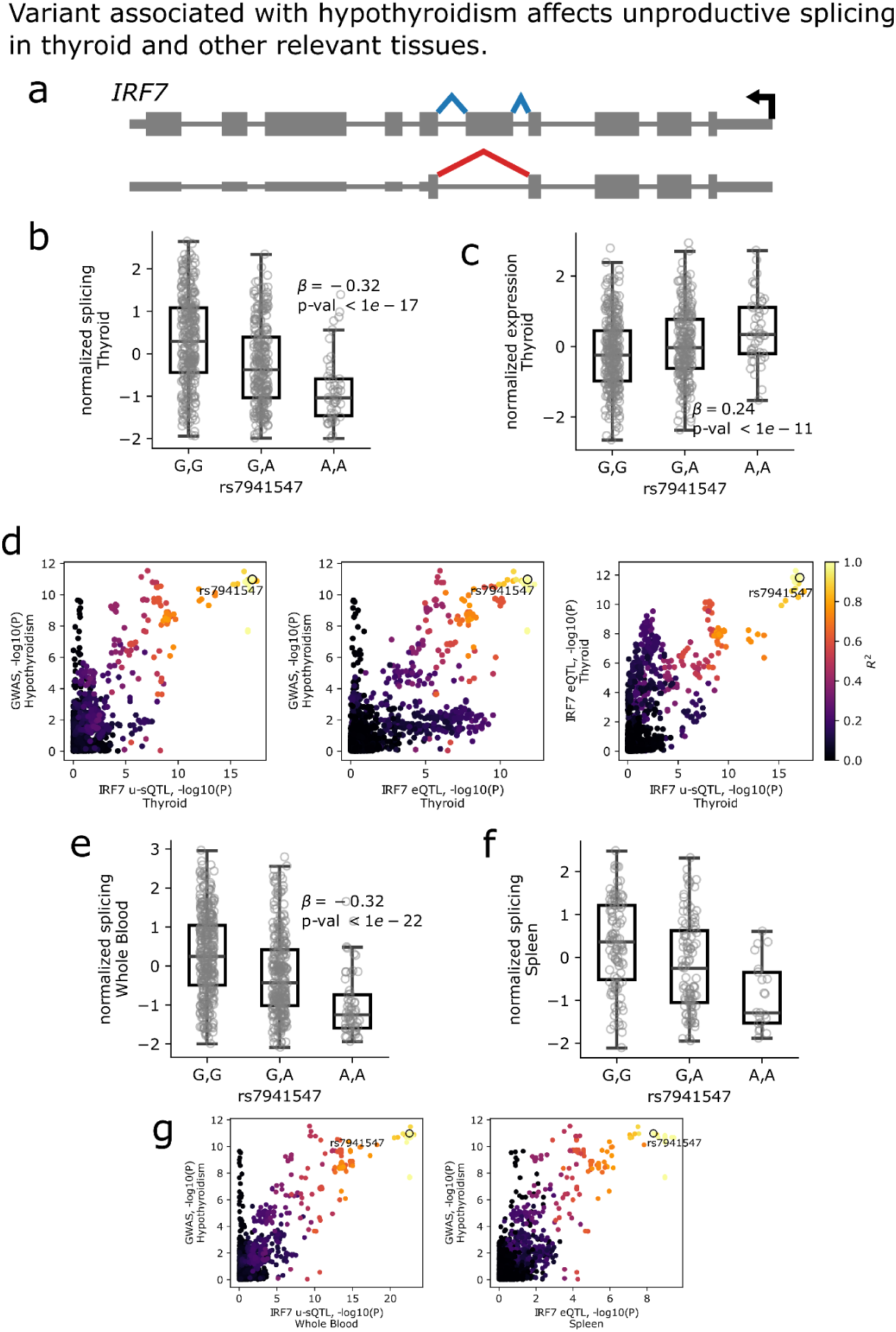
Hypothyroidism variant in *IRF7* is associated with unproductive splicing and gene expression changes. **a)** Productive-unproductive alternative splicing event in *IRF7*. **b)** Variant rs7941547, strongly associated with hypothyroidism, has a strong effect on *IRF7* unproductive splicing in Thyroid tissue. **c)** Variant rs7941547 has a strong effect on *IRF7* expression in a direction consistent with AS-NMD in Thyroid tissue. **d)** LocusCompare plots between Hypothyroidism GWAS signal, u-sQTL effect in IRF7 in Thyroid tissue, and eQTL effect in *IRF7* in Thyroid tissue. **e)** Variant rs7941547 has a strong effect on *IRF7* unproductive splicing in other potentially relevant tissues such as Whole Blood and f) Spleen. **g)** LocusCompare plots between Hypothyroidism GWAS signal, u-sQTL effect in *IRF7* in Whole Blood and Spleen tissue.

**Supplementary Figure 18.**
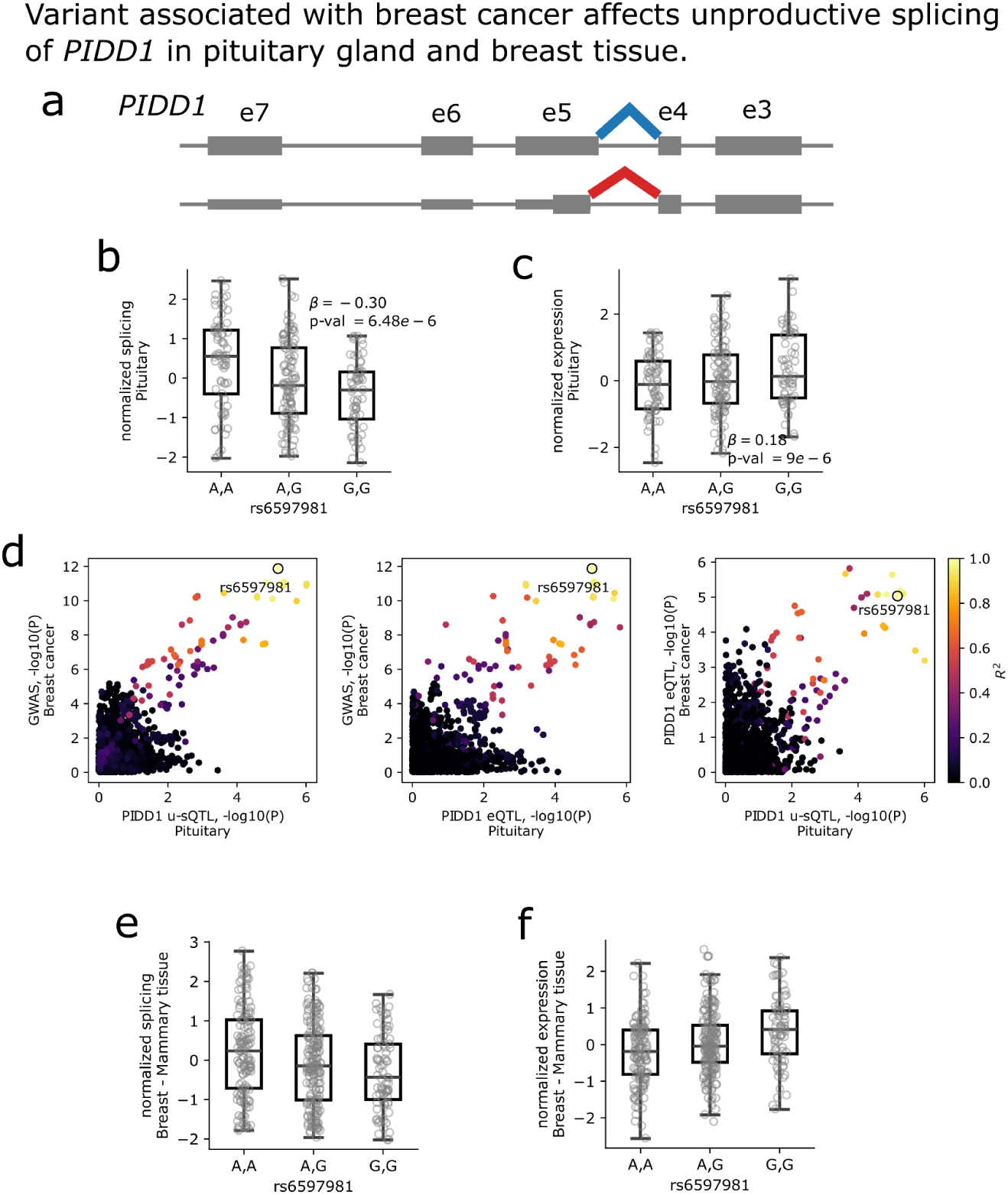
Breast cancer variant in *PIDD1* is associated with unproductive splicing and gene expression changes. **a)** Productive-unproductive alternative splicing event in *PIDD1*. **b)** Variant rs6597981, strongly associated with breast cancer, has a strong effect on *PIDD1* unproductive splicing in Pituitary tissue. **c)** Variant rs6597981 has a strong effect on *PIDD7* expression in a direction consistent with AS-NMD in Pituitary tissue. **d)** LocusCompare plots between Hypothyroidism GWAS signal, u-sQTL effect in *PIDD1* in Pituitary tissue, and eQTL effect in PIDD1 in Pituitary tissue. **e)** Variant rs6597981 has an effect on *PIDD1* unproductive splicing and f) expression in breast mammary tissue. These effects are consistent with the observations in Pituitary tissue, but the u-sQTL effect barely misses the significance threshold.

**Supplementary Figure 19.**
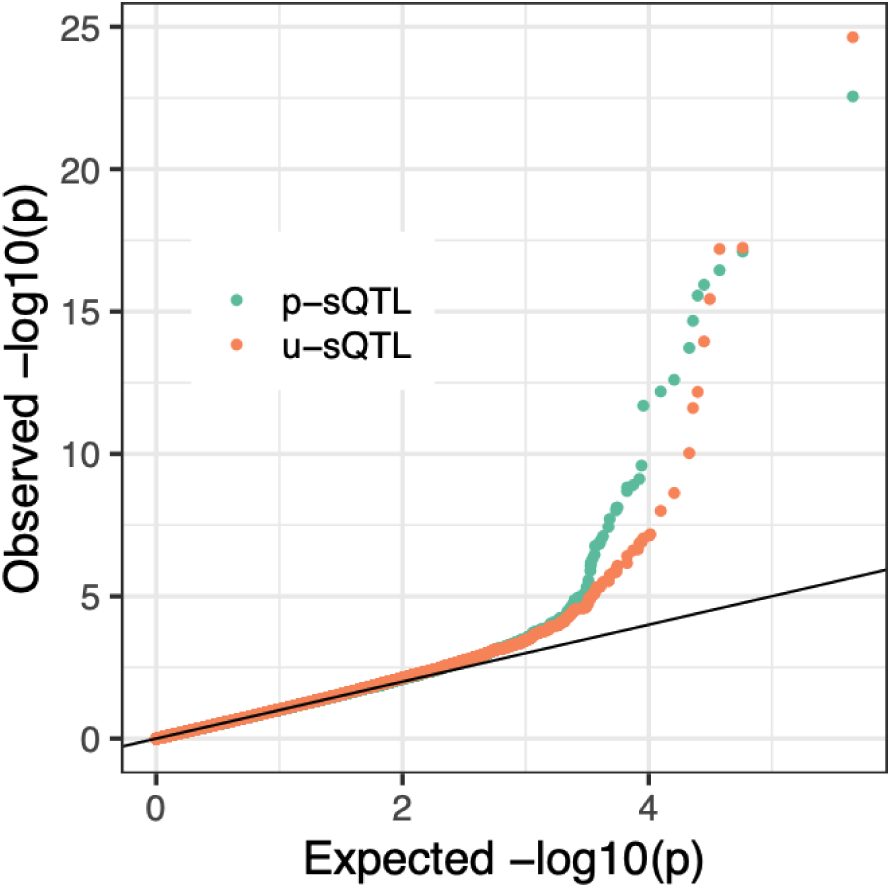
Enrichment of haQTL p-values for u-sQTLs and p-sQTLs. Unproductive splicing QTLs (u-sQTLs) did not exhibit significantly different enrichment in haQTL compared to productive splicing QTLs (p-sQTLs).

**Supplementary Figure 20.**
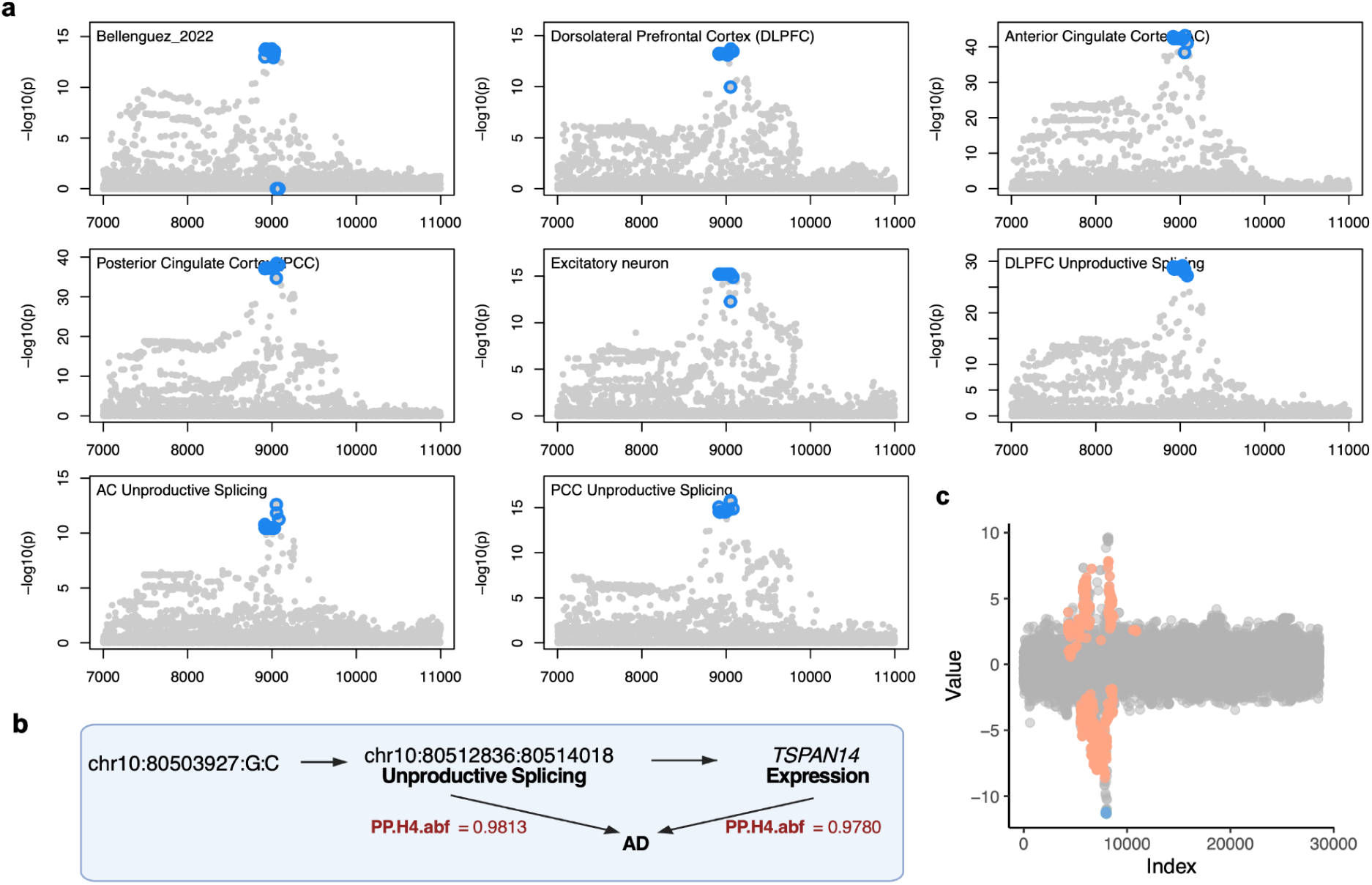
ColocBoost analysis and proposed regulatory model for TSPAN14 unproductive splicing in AD. (a) Colocalization signals are observed among AD risk and unproductive splicing of TSPAN14 in AC, DLPFC, and PCC brain regions (chr10:80512836-80514018), as well as TSPAN14 gene expression and excitatory neuron (Exc) expression. (b) A proposed regulatory model suggests that genetic variants in linkage disequilibrium near chr10:80503927:G:C may drive both unproductive splicing and altered TSPAN14 expression. Colocalization posterior probabilities (PP.H4.abf in SuSiE-COLOC) with AD risk loci are shown in red text. (c) Scatter plot illustrating the Z-scores for variants associated with TSPAN14 unproductive splicing event. Variants involved in potential mediation of unproductive splicing to expression in DLPFC are highlighted in orange (FDR < 0.05 but not the reserved direction). Blue dots denote a 95% colocalized confidence set identified by ColocBoost, which overlaps with the fine-mapping 95% credible set identified by SuSiE.

**Supplementary Table 1.**

| Tissue | Total samples | Samples with genotype | p-sQTL clusters | u-sQTLs clusters |
| --- | --- | --- | --- | --- |
| Adipose - Subcutaneous | 663 | 581 | 6490 | 3942 |
| Adipose - Visceral (Omentum) | 541 | 469 | 5307 | 3126 |
| Adrenal Gland | 258 | 233 | 2986 | 1582 |
| Artery - Aorta | 432 | 387 | 4637 | 2522 |
| Artery - Coronary | 240 | 213 | 2595 | 1443 |
| Artery - Tibial | 663 | 584 | 6255 | 3583 |
| Brain - Amygdala | 152 | 129 | 1130 | 579 |
| Brain - Anterior cingulate cortex (BA24) | 176 | 147 | 1634 | 764 |
| Brain - Caudate (basal ganglia) | 246 | 194 | 2196 | 1091 |
| Brain - Cerebellar Hemisphere | 215 | 175 | 2612 | 1618 |
| Brain - Cerebellum | 241 | 209 | 3092 | 1865 |
| Brain - Cortex | 255 | 205 | 2550 | 1343 |
| Brain - Frontal Cortex (BA9) | 209 | 175 | 2030 | 1012 |
| Brain - Hippocampus | 197 | 165 | 1458 | 766 |
| Brain - Hypothalamus | 202 | 170 | 1782 | 918 |
| Brain - Nucleus accumbens (basal ganglia) | 246 | 202 | 2186 | 1158 |
| Brain - Putamen (basal ganglia) | 205 | 170 | 1723 | 831 |
| Brain - Spinal cord (cervical c-1) | 159 | 126 | 1358 | 721 |
| Brain - Substantianigra | 139 | 114 | 995 | 493 |
| Breast - Mammary Tissue | 459 | 396 | 4745 | 2963 |
| Cells - Cultured fibroblasts | 504 | 483 | 6723 | 3168 |
| Cells - EBV-transformed lymphocytes | 174 | 147 | 2785 | 1379 |
| Colon - Sigmoid | 373 | 318 | 3954 | 2293 |
| Colon - Transverse | 406 | 368 | 4345 | 2551 |
| Esophagus - Gastroesophageal Junction | 375 | 330 | 4180 | 2330 |
| Esophagus - Mucosa | 555 | 497 | 5417 | 2968 |
| Esophagus - Muscularis | 515 | 465 | 5419 | 3005 |
| Heart - Atrial Appendage | 429 | 372 | 4125 | 2158 |
| Heart - Left Ventricle | 432 | 386 | 3250 | 1554 |
| Kidney - Cortex | 85 | 73 | 629 | 332 |
| Liver | 226 | 208 | 1766 | 1043 |
| Lung | 578 | 515 | 5911 | 3598 |
| Minor Salivary Gland | 162 | 144 | 1998 | 1119 |
| Muscle - Skeletal | 803 | 706 | 6146 | 3024 |
| Nerve - Tibial | 619 | 532 | 6532 | 4131 |
| Ovary | 180 | 167 | 2202 | 1421 |
| Pancreas | 328 | 305 | 2724 | 1595 |
| Pituitary | 283 | 237 | 3346 | 2057 |
| Prostate | 245 | 221 | 2679 | 1800 |
| Skin - Not Sun Exposed (Suprapubic) | 604 | 517 | 5639 | 3417 |
| Skin - Sun Exposed (Lower leg) | 701 | 605 | 6353 | 3782 |
| Small Intestine - Terminal Ileum | 187 | 174 | 2281 | 1368 |
| Spleen | 241 | 227 | 3082 | 1870 |
| Stomach | 359 | 324 | 3261 | 1894 |
| Testis | 361 | 322 | 7107 | 4589 |
| Thyroid | 653 | 574 | 6985 | 4169 |
| Uterus | 142 | 129 | 1814 | 1072 |
| Vagina | 156 | 141 | 1746 | 1082 |
| Whole Blood | 755 | 670 | 4080 | 2413 |
| **All tissues** | **17,329** | **15,201** | **9387** | **5107** |

**Supplementary Table 2.**

| **Term** | **Overlap** | **P-value** | **Adjusted P-value** | **Odds Ratio** | **Cluster** |
| --- | --- | --- | --- | --- | --- |
| GOBP RESPONSE TO IONIZING RADIATION | 28 / 144 | 1.79e-07 | 1.88e-04 | 3.55 | Groups III/IV/V (Other tissues) |
| GOBP CELLULAR RESPONSE TO DNA DAMAGE STIMULUS | 108 / 868 | 5.97e-11 | 3.13e-07 | 2.14 | Groups III/IV/V (Other tissues) |
| GOBP DNA REPAIR | 75 / 584 | 1.52e-08 | 3.30e-05 | 2.19 | Groups III/IV/V (Other tissues) |
| GOBP MONOSACCHARIDE METABOLIC PROCESS | 29 / 256 | 1.91e-06 | 2.11e-03 | 2.96 | Groups I/II (Brain) |
| GOBP NCRNA METABOLIC PROCESS | 55 / 613 | 1.95e-07 | 8.56e-04 | 2.31 | Groups I/II (Brain) |
| GOBP NCRNA PROCESSING | 42 / 433 | 7.49e-07 | 1.10e-03 | 2.50 | Groups I/II (Brain) |
| GOBP DNA INTEGRITY CHECKPOINT SIGNALING | 23 / 132 | 1.49e-05 | 4.27e-03 | 3.10 | Groups III/IV/V (Other tissues) |
| GOBP CARBOHYDRATE METABOLIC PROCESS | 53 / 594 | 3.89e-07 | 8.56e-04 | 2.29 | Groups I/II (Brain) |
| GOBP DNA METABOLIC PROCESS | 115 / 1043 | 1.89e-08 | 3.30e-05 | 1.86 | Groups III/IV/V (Other tissues) |
| GOBP RNA SPLICING | 59 / 463 | 6.93e-07 | 5.14e-04 | 2.15 | Groups III/IV/V (Other tissues) |
| GOBP CELL CYCLE CHECKPOINT SIGNALING | 29 / 188 | 1.44e-05 | 4.27e-03 | 2.68 | Groups III/IV/V (Other tissues) |
| GOBP DOUBLE STRAND BREAK REPAIR | 42 / 303 | 3.38e-06 | 1.61e-03 | 2.36 | Groups III/IV/V (Other tissues) |
| GOBP HISTONE MODIFICATION | 60 / 478 | 9.27e-07 | 5.40e-04 | 2.12 | Groups III/IV/V<br>(Other tissues) |
| GOBP ORGANELLE ASSEMBLY | 108 / 993 | 1.07e-07 | 1.40e-04 | 1.82 | Groups III/IV/V<br>(Other tissues) |
| GOBP POSITIVE REGULATION OF<br>RESPONSE TO DNA DAMAGE<br>STIMULUS | 26 / 167 | 3.33e-05 | 7.29e-03 | 2.71 | Groups III/IV/V<br>(Other tissues) |
| GOBP SIGNAL TRANSDUCTION IN<br>RESPONSE TO DNA DAMAGE | 28 / 184 | 2.59e-05 | 6.06e-03 | 2.63 | Groups III/IV/V<br>(Other tissues) |
| GOBP CYTOSKELETON<br>ORGANIZATION | 53 / 1509 | 2.90e-06 | 8.76e-03 | 2.16 | Group VI (Testis) |
| GOBP MRNA METABOLIC<br>PROCESS | 86 / 767 | 6.40e-07 | 5.14e-04 | 1.87 | Groups III/IV/V<br>(Other tissues) |
| GOBP RNA LOCALIZATION | 29 / 197 | 3.52e-05 | 7.40e-03 | 2.53 | Groups III/IV/V<br>(Other tissues) |
| GOBP REGULATION OF DNA<br>METABOLIC PROCESS | 64 / 539 | 2.75e-06 | 1.45e-03 | 1.99 | Groups III/IV/V<br>(Other tissues) |
| GOBP DNA RECOMBINATION | 43 / 328 | 1.09e-05 | 3.90e-03 | 2.21 | Groups III/IV/V<br>(Other tissues) |
| GOBP NCRNA METABOLIC<br>PROCESS | 70 / 613 | 3.82e-06 | 1.67e-03 | 1.90 | Groups III/IV/V<br>(Other tissues) |
| GOBP CELLULAR RESPONSE TO<br>DNA DAMAGE STIMULUS | 66 / 868 | 4.46e-06 | 3.93e-03 | 1.93 | Groups I/II (Brain) |
| GOBP CILIUM ORGANIZATION | 51 / 421 | 1.63e-05 | 4.27e-03 | 2.02 | Groups III/IV/V<br>(Other tissues) |
| GOBP POSITIVE REGULATION OF<br>DNA METABOLIC PROCESS | 39 / 303 | 4.17e-05 | 8.37e-03 | 2.17 | Groups III/IV/V<br>(Other tissues) |
| GOBP CELLULAR RESPONSE TO<br>STRESS | 182 / 1973 | 7.83e-07 | 5.14e-04 | 1.53 | Groups III/IV/V<br>(Other tissues) |
| GOBP CELL PROJECTION<br>ASSEMBLY | 68 / 609 | 1.11e-05 | 3.90e-03 | 1.85 | Groups III/IV/V<br>(Other tissues) |
| GOBP MRNA PROCESSING | 57 / 497 | 2.66e-05 | 6.06e-03 | 1.90 | Groups III/IV/V<br>(Other tissues) |

**Supplementary Table 3.**

| **GWAS trait** | **Loci** | **u-sQTL colocate** | **Percentage colocated** |
| --- | --- | --- | --- |
| Age when finished full-time education | 307 | 27 | 8.79% |
| Asthma childhood onset | 109 | 9 | 8.26% |
| Atopic eczema | 70 | 1 | 1.43% |
| Atrial fibrillation | 111 | 4 | 3.60% |
| Basal cell carcinoma | 94 | 7 | 7.45% |
| Bipolar disorder | 65 | 11 | 16.92% |
| Breast cancer | 168 | 12 | 7.14% |
| Chronic obstructive pulmonary disease | 432 | 33 | 7.64% |
| Coronary artery disease | 221 | 22 | 9.95% |
| Crohn's disease | 97 | 6 | 6.19% |
| Dupuytren's disease | 64 | 4 | 6.25% |
| Heart failure | 42 | 5 | 11.90% |
| Hypothyroidism | 151 | 12 | 7.95% |
| Inflammatory bowel disease | 141 | 12 | 8.51% |
| Multiple sclerosis | 76 | 11 | 14.47% |
| Myocardial infarction | 84 | 8 | 9.52% |
| Rheumatoid arthritis | 76 | 0 | 0% |
| Schizophrenia | 202 | 17 | 8.42% |
| Ulcerative colitis | 62 | 5 | 8.06% |
| Visceral adipose tissue measurement | 325 | 30 | 9.23% |
| **Total** | **2897** | **236** | **8.15%** |

**Supplementary Table 4.**

| **chr** | **Gene ID** | **Gene name** |
| --- | --- | --- |
| 1 | ENSG00000076356 | *PLXNA2* |
| 1 | ENSG00000117322 | *CR2* |
| 1 | ENSG00000203710 | *CR1* |
| 2 | ENSG00000011523 | *CEP68* |
| 2 | ENSG00000071051 | *NCK2* |
| 2 | ENSG00000072135 | *PTPN18* |
| 2 | ENSG00000072163 | *LIMS2* |
| 2 | ENSG00000119147 | *ECRG4* |
| 2 | ENSG00000123636 | *BAZ2B* |
| 2 | ENSG00000136710 | *CCDC115* |
| 2 | ENSG00000136717 | *BIN1* |
| 2 | ENSG00000136731 | *UGGT1* |
| 2 | ENSG00000138380 | *CARF* |
| 2 | ENSG00000138442 | *WDR12* |
| 2 | ENSG00000143951 | *WDPCP* |
| 2 | ENSG00000143952 | *VPS54* |
| 2 | ENSG00000143995 | *MEIS1* |
| 2 | ENSG00000144426 | *NBEAL1* |
| 2 | ENSG00000163161 | *ERCC3* |
| 2 | ENSG00000163596 | *ICA1L* |
| 2 | ENSG00000168918 | *INPP5D* |
| 3 | ENSG00000163655 | *GMPS* |
| 3 | ENSG00000169282 | *KCNAB1* |
| 3 | ENSG00000174953 | *DHX36* |
| 4 | ENSG00000087008 | *ACOX3* |
| 4 | ENSG00000109452 | *INPP4B* |
| 4 | ENSG00000109458 | *GAB1* |
| 4 | ENSG00000127415 | *IDUA* |
| 4 | ENSG00000145214 | *DGKQ* |
| 4 | ENSG00000174227 | *PIGG* |
| 4 | ENSG00000215375 | *MYL5* |
| 5 | ENSG00000050748 | *MAPK9* |
| 5 | ENSG00000051596 | *THOC3* |
| 5 | ENSG00000113119 | *TMCO6* |
| 5 | ENSG00000113194 | *FAF2* |
| 5 | ENSG00000113249 | *HAVCR1* |
| 5 | ENSG00000120725 | *SIL1* |
| 5 | ENSG00000127022 | *CANX* |
| 5 | ENSG00000135077 | *HAVCR2* |
| 5 | ENSG00000145715 | *RASA1* |
| 5 | ENSG00000145850 | *TIMD4* |
| 5 | ENSG00000145901 | *TNIP1* |
| 5 | ENSG00000146090 | *RASGEF1C* |
| 5 | ENSG00000146094 | *DOK3* |
| 5 | ENSG00000155508 | *CNOT8* |
| 5 | ENSG00000183258 | *DDX41* |
| 5 | ENSG00000204962 | *PCDHA8* |
| 5 | ENSG00000285476 | *AC139491.7* |
| 6 | ENSG00000146122 | *DAAM2* |
| 6 | ENSG00000249853 | *HS3ST5* |
| 7 | ENSG00000005020 | *SKAP2* |
| 7 | ENSG00000006530 | *AGK* |
| 7 | ENSG00000106261 | *ZKSCAN1* |
| 7 | ENSG00000106327 | *TFR2* |
| 7 | ENSG00000106351 | *AGFG2* |
| 7 | ENSG00000106460 | *TMEM106B* |
| 7 | ENSG00000146648 | *EGFR* |
| 7 | ENSG00000153814 | *JAZF1* |
| 7 | ENSG00000157800 | *SLC37A3* |
| 7 | ENSG00000159784 | *FAM131B* |
| 7 | ENSG00000159840 | *ZYX* |
| 7 | ENSG00000185899 | *TAS2R60* |
| 7 | ENSG00000197037 | *ZSCAN25* |
| 7 | ENSG00000257923 | *CUX1* |
| 8 | ENSG00000012232 | *EXTL3* |
| 8 | ENSG00000070756 | *PABPC1* |
| 8 | ENSG00000104517 | *UBR5* |
| 8 | ENSG00000120885 | *CLU* |
| 8 | ENSG00000120899 | *PTK2B* |
| 8 | ENSG00000168077 | *SCARA3* |
| 8 | ENSG00000174226 | *SNX31* |
| 8 | ENSG00000186106 | *ANKRD46* |
| 8 | ENSG00000186918 | *ZNF395* |
| 10 | ENSG00000048740 | *CELF2* |
| 10 | ENSG00000059573 | *ALDH18A1* |
| 10 | ENSG00000095585 | *BLNK* |
| 10 | ENSG00000095637 | *SORBS1* |
| 10 | ENSG00000107679 | *PLEKHA1* |
| 10 | ENSG00000108219 | *TSPAN14* |
| 10 | ENSG00000122870 | *BICC1* |
| 10 | ENSG00000122873 | *CISD1* |
| 10 | ENSG00000148429 | *USP6NL* |
| 10 | ENSG00000151150 | *ANK3* |
| 10 | ENSG00000151151 | *IPMK* |
| 10 | ENSG00000155229 | *MMS19* |
| 10 | ENSG00000166033 | *HTRA1* |
| 10 | ENSG00000185737 | *NRG3* |
| 11 | ENSG00000030066 | *NUP160* |
| 11 | ENSG00000073921 | *PICALM* |
| 11 | ENSG00000134569 | *LRP4* |
| 11 | ENSG00000137642 | *SORL1* |
| 11 | ENSG00000149196 | *HIKESHI* |
| 11 | ENSG00000149201 | *CCDC81* |
| 11 | ENSG00000150672 | *DLG2* |
| 11 | ENSG00000150687 | *PRSS23* |
| 11 | ENSG00000165915 | *SLC39A13* |
| 11 | ENSG00000166801 | *FAM111A* |
| 11 | ENSG00000172409 | *CLP1* |
| 12 | ENSG00000089060 | *SLC8B1* |
| 12 | ENSG00000089127 | *OAS1* |
| 12 | ENSG00000089169 | *RPH3A* |
| 12 | ENSG00000123066 | *MED13L* |
| 12 | ENSG00000135094 | *SDS* |
| 12 | ENSG00000139405 | *RITA1* |
| 12 | ENSG00000173064 | *HECTD4* |
| 12 | ENSG00000186815 | *TPCN1* |
| 14 | ENSG00000012963 | *UBR7* |
| 14 | ENSG00000066427 | *ATXN3* |
| 14 | ENSG00000100478 | *AP4S1* |
| 14 | ENSG00000100599 | *RIN3* |
| 14 | ENSG00000100605 | *ITPK1* |
| 14 | ENSG00000100883 | *SRP54* |
| 14 | ENSG00000100890 | *PRORP* |
| 14 | ENSG00000133958 | *UNC79* |
| 14 | ENSG00000140090 | *SLC24A4* |
| 14 | ENSG00000165943 | *MOAP1* |
| 15 | ENSG00000081014 | *AP4E1* |
| 15 | ENSG00000138613 | *APH1B* |
| 15 | ENSG00000140455 | *USP3* |
| 15 | ENSG00000157470 | *FAM81A* |
| 15 | ENSG00000180304 | *OAZ2* |
| 16 | ENSG00000064270 | *ATP2C2* |
| 16 | ENSG00000103175 | *WFDC1* |
| 16 | ENSG00000135686 | *KLHL36* |
| 16 | ENSG00000140943 | *MBTPS1* |
| 16 | ENSG00000197943 | *PLCG2* |
| 17 | ENSG00000011143 | *MKS1* |
| 17 | ENSG00000030582 | *GRN* |
| 17 | ENSG00000067596 | *DHX8* |
| 17 | ENSG00000073969 | *NSF* |
| 17 | ENSG00000178852 | *EFCAB13* |
| 18 | ENSG00000101695 | *RNF125* |
| 18 | ENSG00000118276 | *B4GALT6* |
| 18 | ENSG00000134758 | *RNF138* |
| 18 | ENSG00000141441 | *GAREM1* |
| 19 | ENSG00000007047 | *MARK4* |
| 19 | ENSG00000010310 | *GIPR* |
| 19 | ENSG00000011304 | *PTBP1* |
| 19 | ENSG00000011422 | *PLAUR* |
| 19 | ENSG00000011478 | *QPCTL* |
| 19 | ENSG00000011485 | *PPP5C* |
| 19 | ENSG00000012061 | *ERCC1* |
| 19 | ENSG00000042753 | *AP2S1* |
| 19 | ENSG00000064687 | *ABCA7* |
| 19 | ENSG00000065000 | *AP3D1* |
| 19 | ENSG00000065268 | *WDR18* |
| 19 | ENSG00000073050 | *XRCC1* |
| 19 | ENSG00000076928 | *ARHGEF1* |
| 19 | ENSG00000079313 | *REXO1* |
| 19 | ENSG00000079385 | *CEACAM1* |
| 19 | ENSG00000079432 | *CIC* |
| 19 | ENSG00000089847 | *ANKRD24* |
| 19 | ENSG00000090372 | *STRN4* |
| 19 | ENSG00000090554 | *FLT3LG* |
| 19 | ENSG00000099804 | *CDC34* |
| 19 | ENSG00000099821 | *POLRMT* |
| 19 | ENSG00000099864 | *PALM* |
| 19 | ENSG00000104783 | *KCNN4* |
| 19 | ENSG00000104852 | *SNRNP70* |
| 19 | ENSG00000104853 | *CLPTM1* |
| 19 | ENSG00000104859 | *CLASRP* |
| 19 | ENSG00000104866 | *PPP1R37* |
| 19 | ENSG00000104884 | *ERCC2* |
| 19 | ENSG00000104885 | *DOT1L* |
| 19 | ENSG00000104936 | *DMPK* |
| 19 | ENSG00000104967 | *NOVA2* |
| 19 | ENSG00000104973 | *MED25* |
| 19 | ENSG00000105255 | *FSD1* |
| 19 | ENSG00000105281 | *SLC1A5* |
| 19 | ENSG00000105287 | *PRKD2* |
| 19 | ENSG00000105321 | *CCDC9* |
| 19 | ENSG00000105372 | *RPS19* |
| 19 | ENSG00000105383 | *CD33* |
| 19 | ENSG00000105402 | *NAPA* |
| 19 | ENSG00000105426 | *PTPRS* |
| 19 | ENSG00000105429 | *MEGF8* |
| 19 | ENSG00000105486 | *LIG1* |
| 19 | ENSG00000105497 | *ZNF175* |
| 19 | ENSG00000105501 | *SIGLEC5* |
| 19 | ENSG00000105732 | *ZNF574* |
| 19 | ENSG00000105737 | *GRIK5* |
| 19 | ENSG00000115266 | *APC2* |
| 19 | ENSG00000118162 | *KPTN* |
| 19 | ENSG00000121289 | *CEP89* |
| 19 | ENSG00000124440 | *HIF3A* |
| 19 | ENSG00000124449 | *IRGC* |
| 19 | ENSG00000124459 | *ZNF45* |
| 19 | ENSG00000125743 | *SNRPD2* |
| 19 | ENSG00000125744 | *RTN2* |
| 19 | ENSG00000125746 | *EML2* |
| 19 | ENSG00000125755 | *SYMPK* |
| 19 | ENSG00000129932 | *DOHH* |
| 19 | ENSG00000130005 | *GAMT* |
| 19 | ENSG00000130202 | *NECTIN2* |
| 19 | ENSG00000130203 | *APOE* |
| 19 | ENSG00000130208 | *APOC1* |
| 19 | ENSG00000130751 | *NPAS1* |
| 19 | ENSG00000131115 | *ZNF227* |
| 19 | ENSG00000131116 | *ZNF428* |
| 19 | ENSG00000141873 | *SLC39A3* |
| 19 | ENSG00000141905 | *NFIC* |
| 19 | ENSG00000142002 | *DPP9* |
| 19 | ENSG00000142530 | *FAM71E1* |
| 19 | ENSG00000159917 | *ZNF235* |
| 19 | ENSG00000160014 | *CALM3* |
| 19 | ENSG00000160570 | *DEDD2* |
| 19 | ENSG00000161091 | *MFSD12* |
| 19 | ENSG00000161249 | *DMKN* |
| 19 | ENSG00000167470 | *MIDN* |
| 19 | ENSG00000167637 | *ZNF283* |
| 19 | ENSG00000167671 | *UBXN6* |
| 19 | ENSG00000167680 | *SEMA6B* |
| 19 | ENSG00000172006 | *ZNF554* |
| 19 | ENSG00000174917 | *MICOS13* |
| 19 | ENSG00000176490 | *DIRAS1* |
| 19 | ENSG00000177051 | *FBXO46* |
| 19 | ENSG00000178150 | *ZNF114* |
| 19 | ENSG00000178386 | *ZNF223* |
| 19 | ENSG00000179954 | *SSC5D* |
| 19 | ENSG00000181027 | *FKRP* |
| 19 | ENSG00000182087 | *TMEM259* |
| 19 | ENSG00000185800 | *DMWD* |
| 19 | ENSG00000186026 | *ZNF284* |
| 19 | ENSG00000187116 | *LILRA5* |
| 19 | ENSG00000187244 | *BCAM* |
| 19 | ENSG00000188624 | *IGFL3* |
| 19 | ENSG00000197380 | *DACT3* |
| 19 | ENSG00000197405 | *C5AR1* |
| 19 | ENSG00000214456 | *PLIN5* |
| 19 | ENSG00000224916 | *APOC4-APOC2* |
| 19 | ENSG00000234465 | *PINLYP* |
| 19 | ENSG00000234906 | *APOC2* |
| 19 | ENSG00000256294 | *ZNF225* |
| 19 | ENSG00000267001 | *AC006538.2* |
| 19 | ENSG00000267022 | *AC067968.1* |
| 19 | ENSG00000267385 | *AC011498.4* |
| 19 | ENSG00000267467 | *APOC4* |
| 19 | ENSG00000267508 | *ZNF285* |
| 19 | ENSG00000267680 | *ZNF224* |
| 19 | ENSG00000268361 | *L34079.1* |
| 19 | ENSG00000269469 | *AC010619.1* |
| 19 | ENSG00000278318 | *ZNF229* |
| 20 | ENSG00000087510 | *TFAP2C* |
| 20 | ENSG00000087589 | *CASS4* |

**Supplementary Table 5.**

| Gene ID | Gene name | Brain regions | PTWAS_PVAL | Colocalization |
| --- | --- | --- | --- | --- |
| ENSG00000073921 | *PICALM* | AC,DLPFC,PCC | 3.75E-30 | eQTL & sQTL & GWAS |
| ENSG00000130208 | *APOC1* | AC,DLPFC,PCC | 1.02E-24 | eQTL & GWAS |
| ENSG00000127415 | *IDUA* | AC,DLPFC,PCC | 4.41E-19 | No Colocalization |
| ENSG00000108219 | *TSPAN14* | AC,DLPFC,PCC | 1.77E-14 | eQTL & sQTL & GWAS |
| ENSG00000087589 | *CASS4* | AC | 1.39E-13 | eQTL & sQTL & GWAS |
| ENSG00000072135 | *PTPN18* | AC,DLPFC,PCC | 4.70E-13 | No Colocalization |
| ENSG00000165915 | *SLC39A13* | DLPFC | 1.11E-08 | eQTL & GWAS |
| ENSG00000087008 | *ACOX3* | AC,DLPFC | 3.45E-08 | No Colocalization |
| ENSG00000163596 | *ICA1L* | AC,DLPFC,PCC | 3.23E-07 | No Colocalization |
| ENSG00000104885 | *DOT1L* | AC,DLPFC,PCC | 6.28E-07 | No Colocalization |
| ENSG00000138442 | *WDR12* | AC,PCC | 8.83E-07 | No Colocalization |
| ENSG00000136717 | *BIN1* | AC,DLPFC,PCC | 1.09E-06 | eQTL & GWAS |
| ENSG00000167671 | *UBXN6* | AC,DLPFC | 1.87E-06 | No Colocalization |
| ENSG00000073969 | *NSF* | AC | 5.94E-06 | No Colocalization |
| ENSG00000064687 | *ABCA7* | AC,DLPFC | 1.50E-05 | No Colocalization |
| ENSG00000104884 | *ERCC2* | AC,DLPFC,PCC | 0.00014341 | eQTL & GWAS |
| ENSG00000141441 | *GAREM1* | AC,DLPFC | 0.00038832 | No Colocalization |

